# SorCS2 controls functional expression of amino acid transporter EAAT3 to protect neurons from oxidative stress and epilepsy-induced pathology

**DOI:** 10.1101/426734

**Authors:** Anna R. Malik, Kinga Szydlowska, Karolina Nizinska, Antonino Asaro, Erwin A. van Vliet, Oliver Popp, Gunnar Dittmar, Anders Nykjaer, Katarzyna Lukasiuk, Eleonora Aronica, Thomas E. Willnow

## Abstract

The family of VPS10P domain receptors emerges as central regulator of intracellular protein sorting in neurons with relevance for various brain pathologies. Here, we identified a unique role for the family member SorCS2 in protection of neurons from oxidative stress and from epilepsy-induced cell death. We show that SorCS2 acts as sorting receptor that targets the neuronal amino acid transporter EAAT3 to the plasma membrane to facilitate import of cysteine, required for synthesis of the reactive oxygen species scavenger glutathione. Absence of SorCS2 activity causes aberrant transport of EAAT3 to lysosome for catabolism and impairs cysteine uptake. As a consequence, SorCS2-deficient mice exhibit oxidative brain damage that coincides with enhanced neuronal cell death and increased mortality during epilepsy. Our findings highlight a protective role for SorCS2 in neuronal stress response and provide an explanation for upregulation of the receptor seen in surviving neurons of the human epileptic brain.

## INTRODUCTION

Sorting of cellular proteins to their correct target sites is fundamental for maintaining proper cell functions. Intracellular protein sorting is particularly challenging for neurons, composed of diverse subcellular compartments in cell soma, dendrites, and axons. Not surpsingly, aberrant protein sorting in neurons has therefore been recognized as underling cause of many devastating brain pathologies (Wang et al., 2013). In recent years, the functional characterization of a unique class of intracellular sorting receptors, termed VPS10P domain receptors, has shed light on the molecular mechanisms that govern protein sorting in neurons. VPS10P domain receptors encompass a group of five structurally related type 1-transmembrane proteins expressed in neurons of the central and peripheral nervous systems (Willnow et al., 2008). Gene family members are SORLA, sortilin, as well as SorCS1, −2 and −3. As a common feature, VPS10P domain receptors act as sorting receptors that direct cargo proteins between Golgi, cell surface, and endosomal compartments. One of their ascribed functions is the ability to move signaling receptors to and from the cell surface, providing essential control of neuronal signal reception. Examples of such activity include sorting of neurotrophin receptors by sortilin (Vaegter et al., 2011) and SorCS1 and −3 (Subkhangulova et al., 2018), or of receptors for glia cell line-derived neurotrophic factor by SORLA (Glerup et al., 2013).

Recently, VPS10P domain receptors have received growing attention owing to their relevance for disorders of the human brain. Thus, genetic studies have associated genes encoding VPS10P domain receptors with psychiatric diseases, including *SORCS1* with attention deficit hyperactivity disorder (ADHD, Lionel et al., 2011) and *SORCS2* with bipolar disorder (Baum et al., 2008; Ollila et al., 2009), schizophrenia (Christoforou et al., 2011) and ADHD (Alemany et al., 2015). Also, epidemiological studies have associated SORLA and sortilin with age-related dementias, including Alzheimer’s disease (Lambert et al., 2013; Rogaeva et al., 2007). Although the exact molecular mechanisms whereby VPS10P domain receptors may impact brain diseases are still not fully understood, faulty protein transport has been identified as one of the underlying reasons, as shown for SORLA as a sorting receptor for the amyloid precursor protein in Alzheimer’s disease (Andersen et al., 2005; Caglayan et al., 2014; Offe et al., 2006; Schmidt et al., 2012).

As well as by genetic association, VPS10P domain receptors have also been implicated in brain pathologies by expression studies identifying altered receptor levels in the diseased brains of patients or animal models (Andersen et al., 2005; Finan et al., 2011; Reitz et al., 2013; Saadipour et al., 2013). In fact, expression studies have been particularly instructive as they provided an unbiased approach to identifying hitherto unknown connections between directed protein sorting and brain pathology. With relevance to this study, *Sorcs2* was identified as a gene strongly upregulated in an experimental mouse model of temporal lobe epilepsy (TLE), implicating this receptor in neuropathologies associated with brain seizures (VonDran et al., 2014). However, the underlying molecular mechanism of SorCS2 function, and whether it bears relevance for epilepsy in humans, remained unclear. Combining unbiased proteome screens for novel SorCS2 targets with studies in receptor-deficient mouse models and in patients with TLE, we now identified the unique function of this receptor in trafficking of the neuronal glutamate/cysteine transporter EAAT3. SorCS2-mediated sorting of EAAT3 to the cell surface promotes uptake of cysteine, the precursor to produce the scavenger for reactive oxygen species glutathione, and it protects neurons from oxidative stress induced by seizures. Jointly, these findings have uncovered the significance of directed protein sorting for control of neuronal stress response and as neuroprotective pathway in epilepsy.

## RESULTS

### SorCS2 is upregulated in the epileptic human brain and plays a protective role in an experimental model of TLE

Epilepsy is a complex chronic brain disorder characterized by seizures, leading to circuit reorganization, neuronal cell loss, as well as other structural and functional brain abnormalities. Pathological features of the human disease can be recapitulated in mouse models of epilepsy involving the administration of chemoconvulsants, such as pilocarpine, kainic acid, or pentylenetetrazol (PTZ). With relevance to this study, SorCS2 emerged as one of the proteins upregulated in an experimental mouse model of TLE. Specifically, SorCS2 levels increased in neurons of the hippocampus three days after status epilepticus induced by pilocarpine. Increased receptor expression was particularly evident in the hilus and cornus ammonis 2 (CA2) region (VonDran et al., 2014). To verify the relevance of this observation for human brain pathology, we tested the expression pattern of SorCS2 in the healthy and the epileptic human hippocampus with hippocampal sclerosis (HS) (Fig. 1 A). HS is the most common type of neuropathological damage seen in individuals with TLE, characterized by severe neuronal cell loss in the hippocampus. In healthy control tissue, we observed robust SorCS2 expression in neurons of the hippocampus, in agreement with the reported expression pattern of the receptor in the mouse brain (Hermey et al., 2004). In human TLE-HS tissue samples, severe neuronal cell loss was visible in all hippocampal regions with the exception of CA2, which is particularly resistant to epilepsy-driven degeneration (Steve et al., 2014). Strikingly, surviving neurons in the CA2 region of human epilepsy samples showed a substantial increase in SorCS2 level, suggesting a potential role for this receptor in preventing neuron loss in the course of epilepsy (Fig. 1 A).

**Figure 1.**
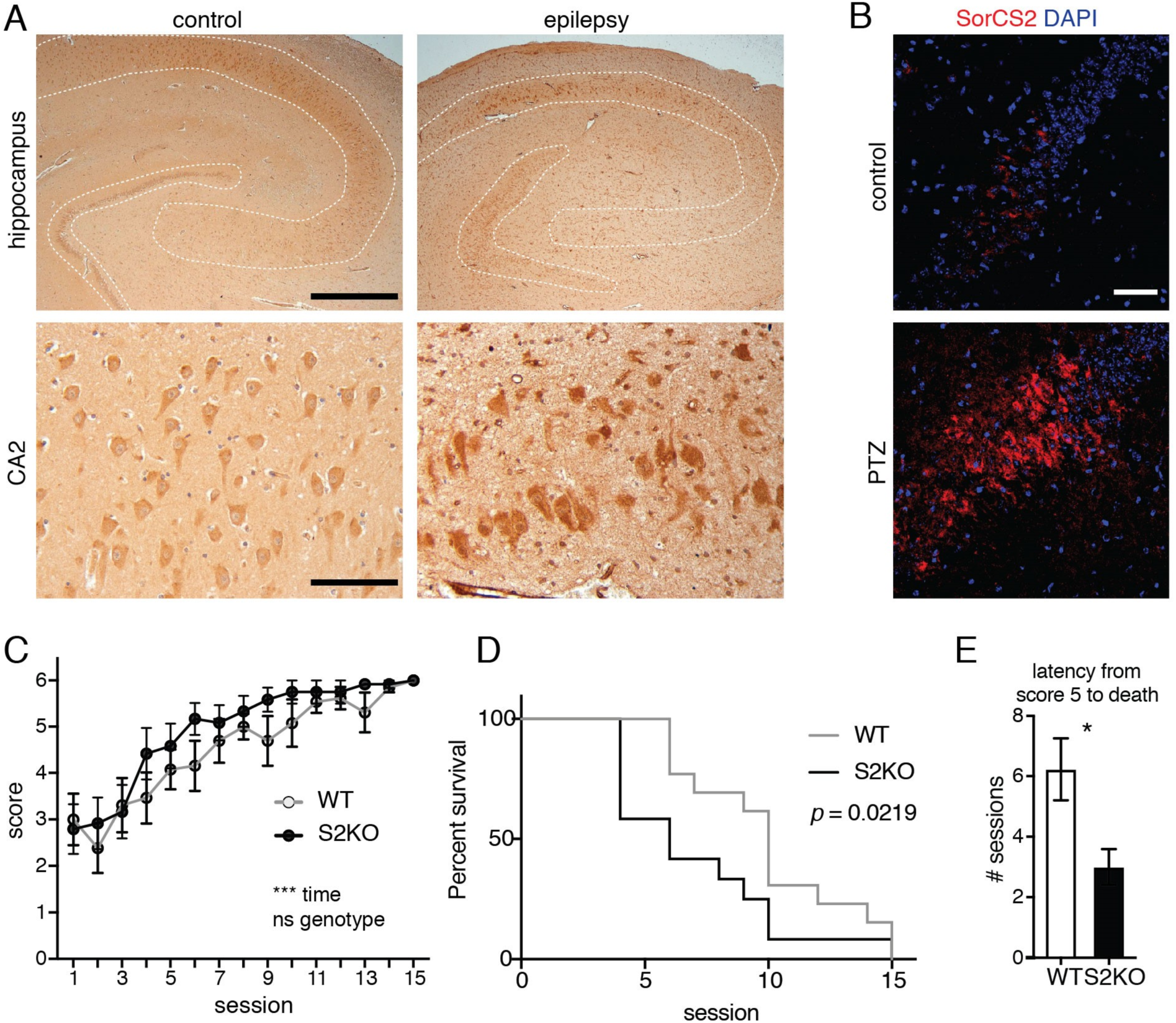
SorCS2 is upregulated in the epileptic human brain and plays a protective role in an experimental model of temporal lobe epilepsy (TLE). (A) SorCS2 immunostaining in the epileptic human hippocampus. Left panels – human control hippocampus, right panels – hippocampus of a patient with TLE and hippocampal sclerosis. Neuronal cell loss is evident in the hippocampal CA and DG (indicted by white dotted lines) of the epileptic as compared to the control brain tissue. SorCS2 expression is visible in all regions of the healthy and the epileptic hippocampus, but massively increases in surviving neurons, in particular in the CA2 region of the epileptic brain. Scale bar: overview, 1 mm; CA2 region, 100 μm. See also Table S1. (B) SorCS2 immunostaining (red) in the CA2 region of the hippocampus of non-treated (control) and PTZ kindled wild-type mice. Sections are counterstained with DAPI. Massive increase in SorCS2 immunoreactivity is seen in the CA2 region of the epileptic brain. Scale bar: 50 μm. (C-E) Results of the PTZ kindling experiments. WT and S2KO mice were injected with PTZ three times a week and their seizures scored (from 0 - no seizure, to 5; 6 – death) as detailed in the method section. n=12–13 mice per genotype. (C) Seizure scores in the function of time (sessions) in the PTZ kindling experiment. Mean value ± SEM; ***, *p*<0.001 (two-way repeated measures ANOVA). (D) Survival curve of WT and S2KO mice in course of the PTZ kindling experiment, *p* = 0.0219 in Gehan-Breslow-Wilcoxon test for comparison of survival curves. (E) Latency (number of sessions) from score 5 to score 6 (death) in course of the PTZ kindling experiment for WT and S2KO mice. Mean value ± SEM; *, *p*<0.05 (unpaired t-test).

To further investigate possible SorCS2 functions in epilepsy, we turned to a PTZ-induced kindling paradigm applied to mice either wild-type (WT) or genetically deficient for *Sorcs2* (referred to as S2KO; Glerup et al., 2014). Kindling is a paradigm whereby repeated low-level stimulation leads to a permanent increase in seizure susceptibility and to the development of seizures, which intensify over time. In this chronic epilepsy model, mice were subjected to a sub-convulsive dose of PTZ three times a week for 5 weeks (15 sessions in total). During the course of the treatment, the convulsive behavior was video-recorded and the animals were scored for the severity of seizures (0 to 5; 6 in case of death; Becker et al., 1992). Over time, mice repeatedly challenged with this pro-convulsive drug develop convulsions of increasing severity. As was the case for the epileptic human brain, we observed substantially increased levels of SorCS2 protein in the CA2 region of the hippocampus in PTZ-kindled WT mice using immunohistochemistry (Fig. 1 B). Overall, S2KO mice presented a kindling phenotype that was comparable to that of WT animals (Fig. 1 C). Also, seizures resulting from the first administration of PTZ, reflecting the acute response to the stimulus, did not differ between the SorCS2-deficient and control animals (Fig. 1 C). However, we observed a striking difference in the mortality rate of the two genotype groups during the course of the kindling experiment with the mortality rate being higher in S2KO mice as compared to WT controls (Fig. 1 D). Of note, once developing strong seizures (score 5), SorSC2-deficient mice died within the next three sessions on average, whereas the WT controls survived for approximately 6 more sessions (p<0.05; Fig. 1 E). This observation indicated that SorCS2-deficient mice were more prone to developing irreversible pathological changes during kindling that eventually led to their death.

### Loss of SorCS2 aggravates cell death and oxidative damage in a murine model of epilepsy

Irreversible pathological changes in S2KO animals may involve neuron loss, a characteristic epilepsy-associated process with a critical impact on the outcome of seizures (Dingledine et al., 2014). To query potential differences in epilepsy-driven cell death, we assessed the number of apoptotic cells in the hippocampi of WT and S2KO mice by TUNEL staining at the endpoint of the PTZ kindling experiment. In line with our hypothesis, the number of TUNEL-positive cells was significantly higher in the CA2/3 regions of S2KO hippocampi as compared to the WT control tissue (Fig. 2 A-B). Since the kindling phenotype, as well as the extent of seizures after the first PTZ dose, were comparable between the genotypes, we concluded that the aggravated cell death observed in S2KO hippocampi was not due to an overall difference in responsiveness to PTZ. More likely, repeated epileptic seizures may lead to secondary changes in brain functions that were more pronounced in the S2KO as compared to WT brains. Since oxidative stress is an important feature of epilepsy and likely contributes to epilepsy-associated cell death (Shin et al., 2011), we tested the levels of the oxidative stress marker 8-hydroxy-2’-deoxyguanosine (8OHdG) in mouse hippocampi after PTZ-kindling. 8OHdG is one of the predominant forms of free radical-induced oxidative lesions in nuclear and mitochondrial DNA (Valavanidis et al., 2009). Remarkably, 8OHdG immunoreactivity was significantly increased in S2KO hippocampi as compared to WT tissue, both in the dentate gyrus (DG) and the CA region (Fig. 2 C-D). This observation supported our concept that the irreversible brain damage induced in S2KO mice by PTZ kindling was related to increased susceptibility to oxidative stress.

**Figure 2.**
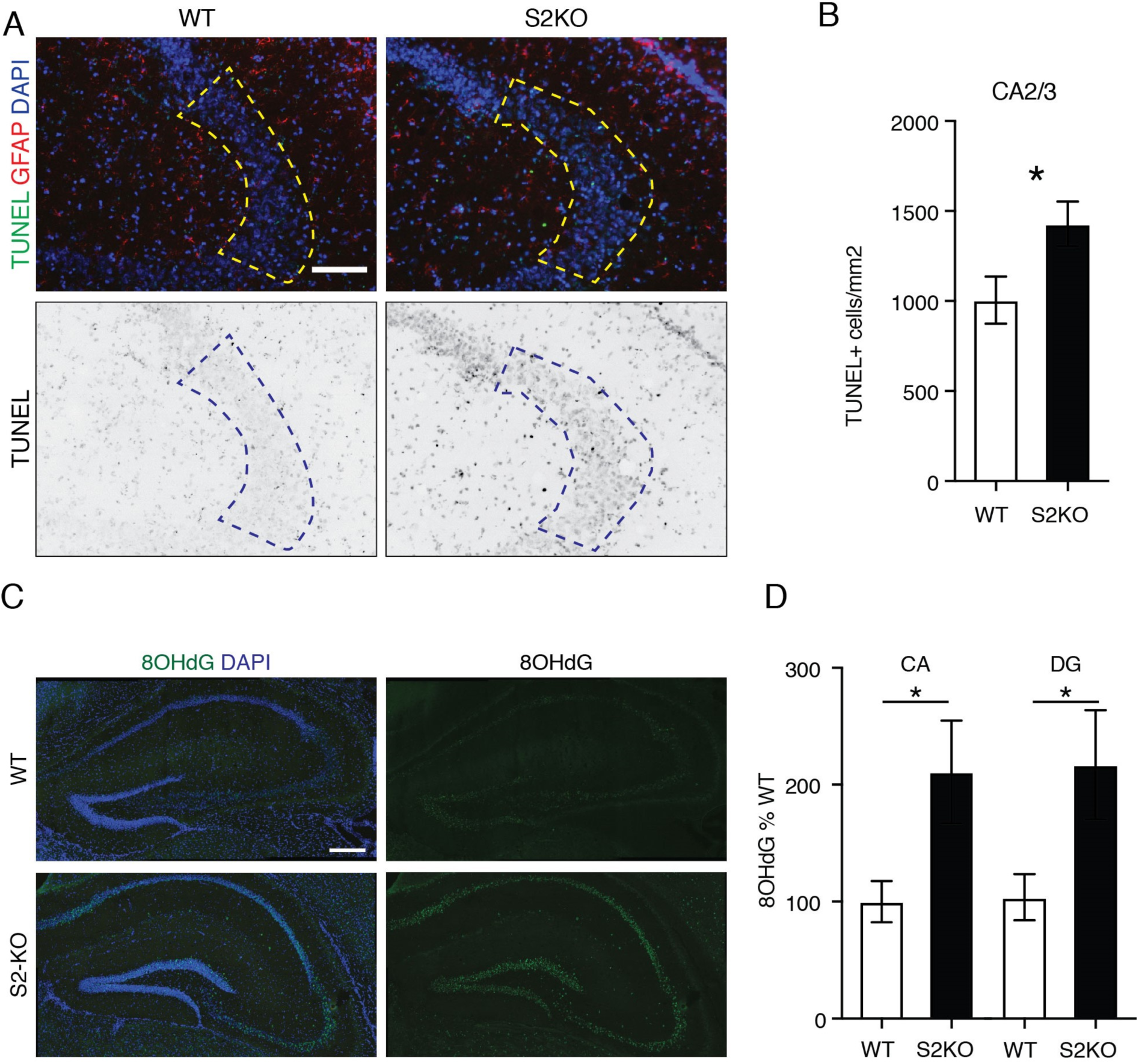
Loss of SorCS2 aggravates cell death and oxidative stress response in a murine model of epilepsy. (A) Representative images of TUNEL staining (green) in the CA2/3 region of the hippocampus of WT or S2KO mice after PTZ kindling (upper panels). Staining for glial fibrillary acid protein (GFAP) marks reactive astrocytes (red). Dashed lines indicate the areas in which the number of TUNEL-positive cells was quantified. For quantification, the green channel was evaluated separately (lower panels, shown in greyscale). Scale bar: 100 μm. (B) Quantification of TUNEL-positive cells in the CA2/3 area of the hippocampus of epileptic WT and S2KO mice (as exemplified in panel A). Mean ± SEM; n=10–11 mice per genotype; *,p< 0.05 (unpaired t-test). (C) Representative images of immunostaining for oxidative stress marker 8OHdG (green) in hippocampi of WT and S2KO mice after PTZ kindling. Cells were counterstained with DAPI (blue). Scale bar: 250 μm. (D) Quantification of 8OHdG immunostaining intensity in the CA1–3 regions (CA) and dentate gyrus (DG) of the WT and S2KO mice after PTZ kindling (as exemplified in panel C). Mean ± SEM; n=12-13 mice per genotype; *, *p*<0.05 (unpaired t-test).

In support of a role for SorCS2 in protection from oxidative stress, we observed that cultured SorCS2 mutant neurons, although in general responding normally to pharmacological stimuli, showed an aberrant stress-related response under certain conditions. In detail, treatment of WT and S2KO primary neurons with the GABA-A receptor antagonist bicuculline resulted in comparable levels of activation of classical mitogen-activated signal transduction pathway, as reflected by similarly induced levels of p-ERK (Fig. 3 A-B). Also, brief exposure to low concentrations of the selective agonist of N-methyl-D-aspartate receptors (NMDA; 20 μM, 5 min) caused activation of the ERK signaling pathway and an increase in the level of p-ERK (Fig. 3 C-D) that was comparable for WT and S2KO neurons. However, stimulation with NMDA also triggered an increase in p-p38 levels that was significantly more pronounced in S2KO as compared to WT neurons (Fig. 3 C, E). In contrast to ERK, p-38 is a component of a signaling cascade that specifically controls cellular response to stress (Roux and Blenis, 2004), including oxidative stress triggered by NMDA (Sengpiel et al., 1998). In summary, our data indicated that SorCS2 is involved in cellular mechanisms providing protection of neurons from oxidative stress.

**Figure 3.**
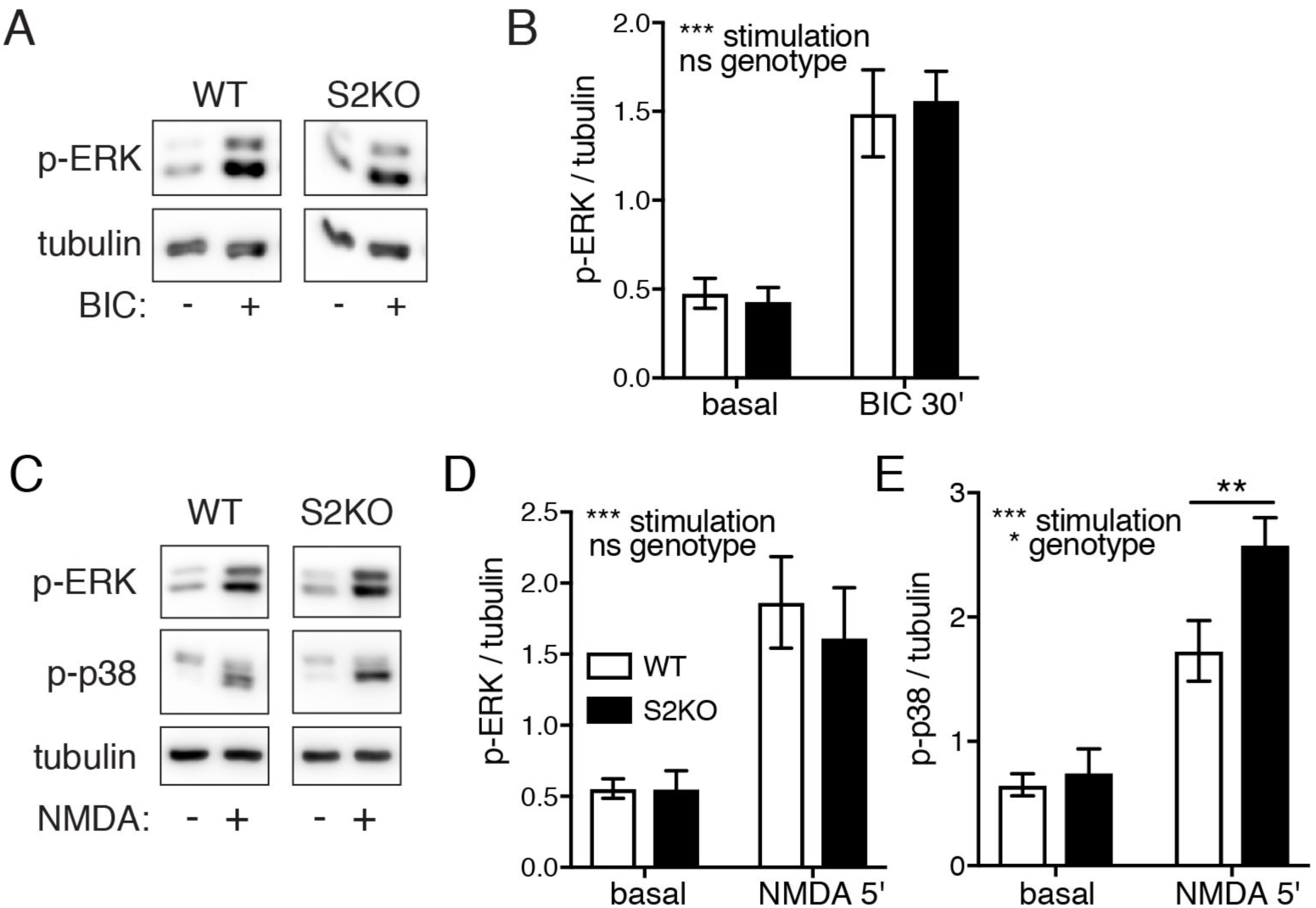
Loss of SorCS2 aggravates stress response in primary neurons. (A) Western blot analysis of p-ERK levels in cell lysates after stimulation of primary WT and S2KO neurons with 20 μM bicuculline (BIC) for 30 minutes. Tubulin is shown as a loading control. (B) Quantification of p-ERK levels in neuronal lysates (normalized to tubulin levels). Mean ± SEM; n=6-7 independent neuronal preparations per genotype; ns, not significant; ***, *p*<0.001 (two-way ANOVA). (C) Western blot analysis of p-ERK and p-p38 levels in cell lysates after stimulation of primary WT and S2KO neurons with 20 μM NMDA for 5 minutes. Tubulin is shown as a loading control. (D-E) Quantification of p-ERK and p-p38 levels in neuronal cell lysates (normalized to tubulin levels). Mean ± SEM; n=6-7 independent neuronal preparations per genotype; ns, not significant; *, *p*<0.05; **, *p*<0.01; ***, *p*<0.001 (two-way ANOVA with Sidak’s multiple comparisons test).

### Neuronal surface proteome is altered in S2KO neurons

To gain a better understanding of SorCS2-dependent cellular processes protecting neurons from oxidative stress, we applied an unbiased proteomics-based approach to identify novel molecular targets of SorCS2 action in the neuronal surface proteome. This approach is based on the established role of VPS10P domain receptors in sorting proteins to and from the neuronal cell surface, and on documented alterations in cargo exposure on the cell surface in cells lacking individual VPS10P domain receptors (Mazella et al., 2010; Rohe et al., 2013). We had applied this strategy successfully before to identify hitherto unknown ligands for gene family members SorCS1 and SorCS3 (Subkhangulova et al., 2018). Here, neuronal cell surface proteins in WT and S2KO primary neurons were biotinylated using cell membrane impermeant compound EZ-Link™Sulfo-NHS-SS-Biotin, and subsequently purified on a streptavidin resin. The resulting neuronal cell surface fractions were subjected to quantitative label-free mass spectrometry analysis (Fig. 4 A). In these experiments, we uncovered multiple proteins with changed abundance in the neuronal cell surface fraction upon loss of SorCS2 (Fig. 4 B, Table 1, and Table S2). Conceptually, identified hits may represent direct targets of SorCS2-dependent trafficking, or proteins altered secondarily as a consequence of faulty neuronal transport processes. Importantly, proteins with decreased cell surface exposure in mutant neurons included SorCS2 itself (Fig. 4 B, Table 1), substantiating the applicabilty of our strategy to identifying alterations in the cell surface proteome of S2KO neurons. Among the hits, we identified cell surface receptors, channels and transporters, as well as proteins implicated in intracellular trafficking processes (Table 1).

**Table 1.**
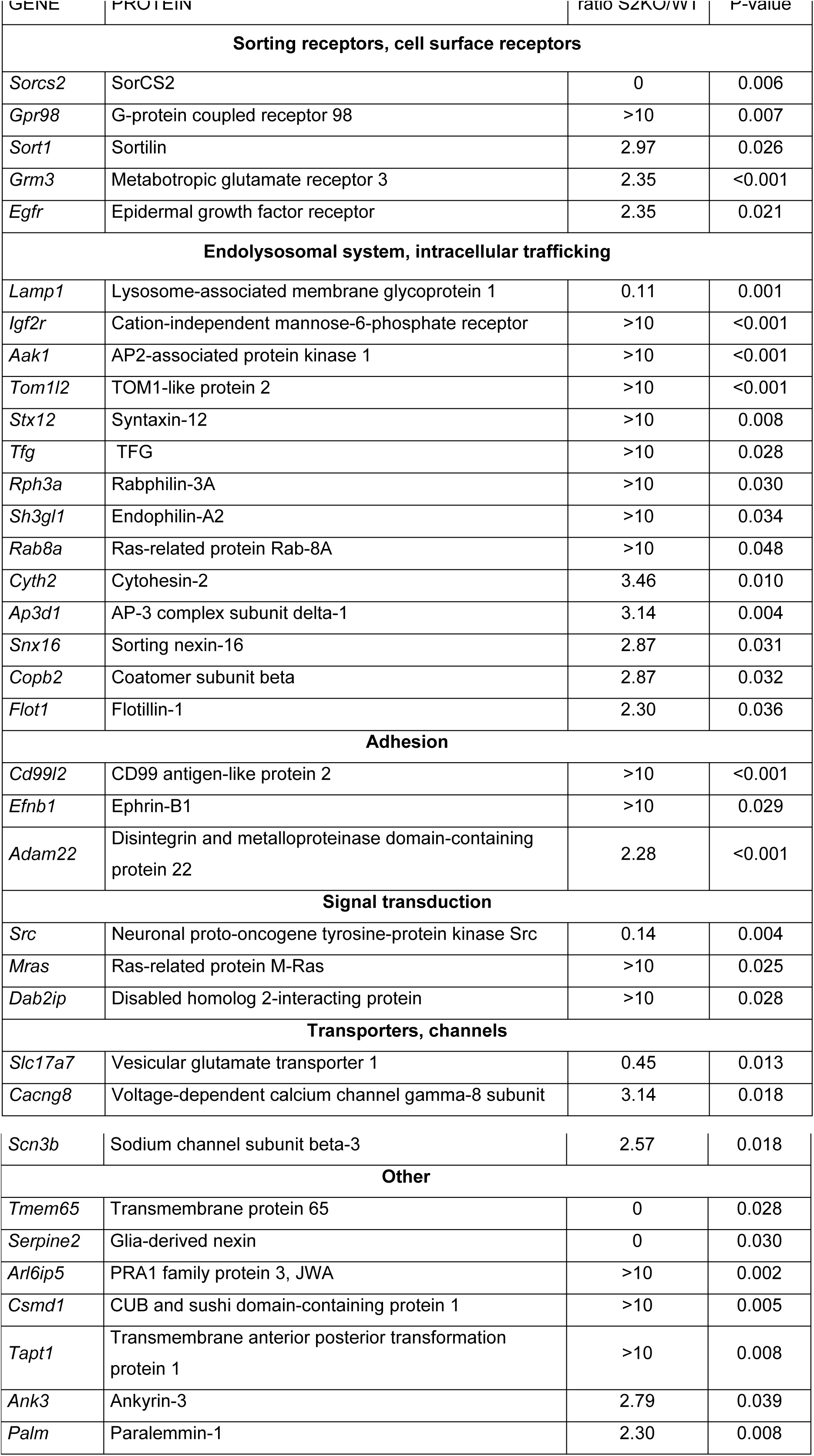
List of selected proteins with altered cell surface exposure in SorCS2-deficient neurons.

**Figure 4.**
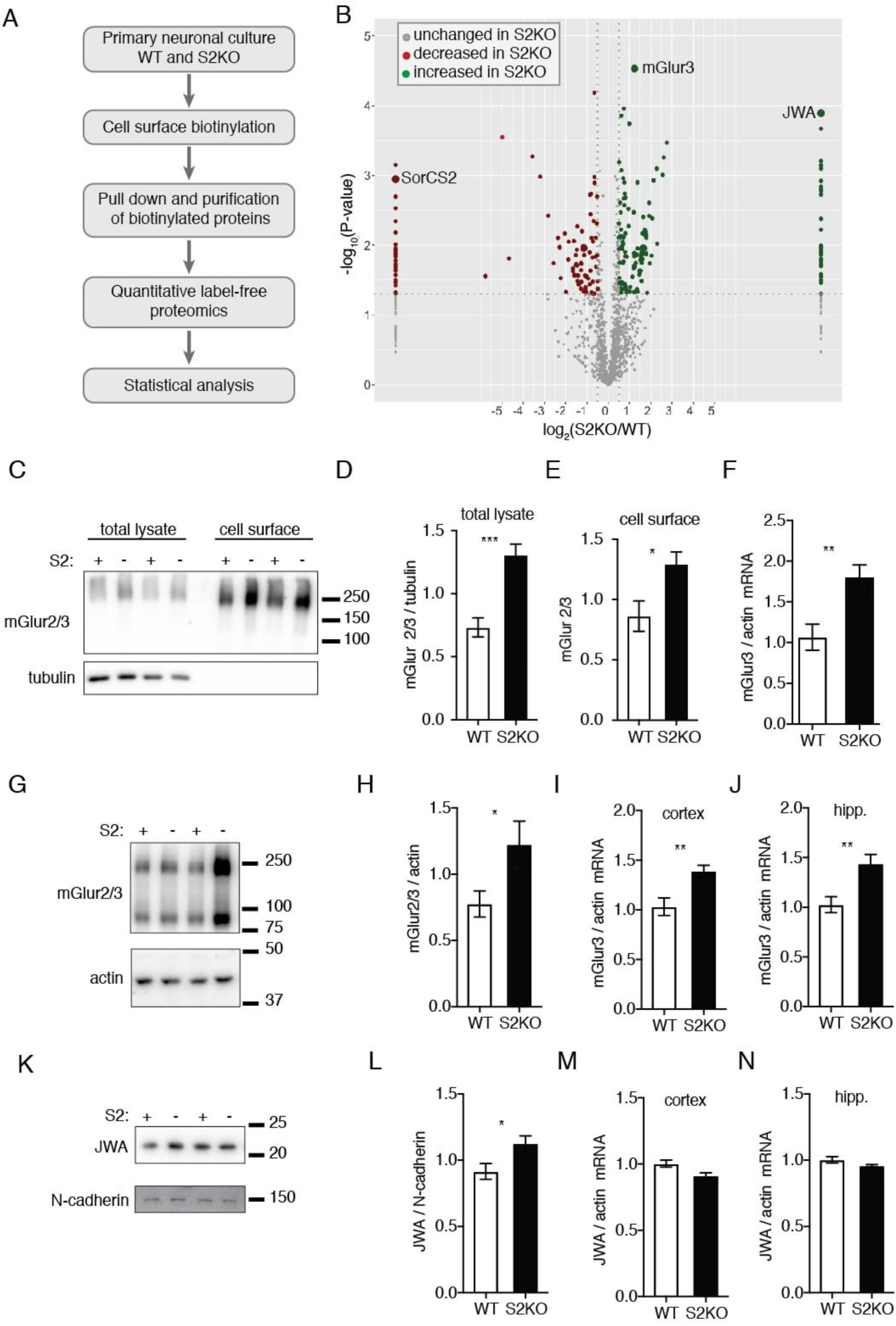
Composition of the cell surface proteome is altered in SorCS2-deficient neurons. (A) Workflow of the cell surface proteome analysis in primary neurons. (B) Results of quantitative label-free proteomics comparing the surface proteomes of WT and S2KO primary neurons. Plot represents −log_10_(*P*-value) and log_2_[relative levels (S2KO/WT)] obtained for each protein. *n=3* biological replicates/group (with each biological replicate run in two technical replicates). See also Table S2. (C) Western blot analysis of total and cell surface levels of mGlur2/3 in WT and S2KO primary neurons. The primary antibody detects both mGlur2 and mGlur3 (multimers, >200kDa, predominant form in cultured neurons). Tubulin is shown as a loading control for the total cell lysate. (D-E) Quantification of mGlur2/3 levels in neuronal lysates (D) and cell surface fractions (E) of WT and S2KO primary neurons (as exemplified in panel C). Mean ± SEM; n=5-6 independent neuronal preparations per genotype; *, *p*<0.05; **, *p*<0.01 (unpaired t-test). (F) mGlur3 mRNA levels in primary WT and S2KO neurons as assessed by quantitative (q) RT-PCR (relative to actin mRNA). Mean ± SEM; n=5-6 independent neuronal preparations per genotype; **, *p*<0.01 (unpaired t-test). (G) Western blot analysis of mGlur2/3 level in P2 brain fraction (crude membranes, see also Figure S2 A). The antibody detects both mGlur2 and mGlur3 (monomers, 90 kDa; multimers, >200 kDa). Actin is shown as a loading control. See also Figure S1. (H) Quantification of mGlur2/3 levels in P2 brain fraction (normalized to actin levels) as exemplified in panel G. Mean ± SEM; n=7 mice per genotype; *, *p*<0.05 (unpaired t-test). (I-J) mGlur3 mRNA levels in brain cortex (i) and hippocampus (hipp.; j) of WT and S2KO mice as assessed by qRT-PCR (relative to actin). Mean ± SEM; n=9 mice per group; **, *p*<0.01 (unpaired t-test). (K) Western blot analysis of JWA levels in brain lysates from WT and S2KO mice. N-cadherin served as loading control. See also Figure S1. (L) Quantification of JWA levels in brain lysate from WT and S2KO mice (normalized to N-cadherin levels) as exemplified in panel k. Mean ± SEM; n=6 mice per genotype; *, p< 0.05 (unpaired t-test). (M-N) JWA mRNA levels in brain cortex (m) and hippocampus (hipp.; n) of WT and S2KO mice as assessed by qRT-PCR (relative to actin). Mean ± SEM; n=9 mice per genotype.

Interestingly, two hits in our proteome screen, namely JWA and mGlur3 were implicated in oxidative stress response. JWA was induced in NIH-3T3 and HELF cells in response to oxidative stress triggered by hydrogen peroxide (Chen et al., 2007) and was implicated in regulation of the synthesis of the reactive oxygen species (ROS) scavenger glutathione (Watabe et al., 2008). Metabotropic glutamate receptors (mGlurs), in addition to their involvement in glutamate-mediated signal transduction, also play a role in protection from oxidative stress. For example, activation of mGlur2 and mGlur3 increased synthesis of glutathione in cultured dorsal root ganglion neurons (Berent-Spillson and Russell, 2007) and prevented the formation of ROS in response to elevated glucose concentrations.

We confirmed an increase in the expression levels of mGlur2/3 in S2KO primary neurons by Western blot (Fig. 4 C-E), substantiating the results from our surface proteome screen. Interestingly, this increase was not limited to the plasma membrane fraction, but was also observed in the total neuronal lysate. We also documented upregulation of mGlur2/3 protein levels in the SorCS2-deficient mouse brain *in vivo* (Fig. 4 G-H). Of note, an increase in mGlur3 expression in S2KO mice was already seen at the transcript levels in both primary neurons (Fig. 4 F) and in brain (Fig. 4 I-J), arguing that this protein was likely not a direct target of SorCS2-mediated protein sorting. An increase in protein level was also substantiated for JWA in brain lysates of SorCS2 mutant mice (Fig. 4 K-L). In this case, mRNA levels remained unchanged between the genotypes (Fig. 4 M-N), suggesting a post-transcriptional mechanism of expression regulation for JWA (as opposed to mGlur3). To query whether JWA or mGlur3 may be direct targets of SorCS2 action in the brain, we tested the ability of both proteins to interact with SorCS2. However, we failed to document co-immunoprecipitation of either protein with SorCS2 (Fig. S1), arguing that alterations in levels of JWA and mGlur3 reflected a secondary consequence of loss of SorCS2.

### SorCS2 co-localizes with the glutamate/cysteine transporter EAAT3

Our proteomics screen and the confirmatory experiments in brain tissue and primary neurons pointed to alterations in glutamatergic and oxidative stress response systems in S2KO neurons. Conceptually, both systems are mechanistically linked as glutamate is not only a neurotransmitter but also a substrate for synthesis of glutathione, the main cellular ROS scavenger. Synthesis of glutathione requires two addtional amino acids, glycine and cysteine. Of note, glutamate and cysteine are imported into neurons by the same amino acid transporter, called excitatory amino acid transporter 3 (EAAT3; or excitatory amino-acid carrier 1, EAAC1). Intriguingly, EAAT3 is functionally linked to hits of our proteomics screen. Thus, JWA is a direct interactor and modulator of EAAT3 trafficking and/or function (Lin et al., 2001; Ruggiero et al., 2008). Also, activation of mGlur2/3 increased levels of EAAT3 in glioma cells (Aronica et al., 2003).

In the mammalian brain, EAAT3 is highly expressed by neurons, where it localizes both to the dendrites and to the cell soma (Holmseth et al., 2012). Within cells, a large fraction of EAAT3 resides in the endosomal compartment, particularly in Rab11-positive recycling endosomes that play a crucial role in EAAT3 intracellular sorting and targeting to the plasma membrane (González et al., 2007; Su et al., 2016). To test possible colocalization of SorCS2 with EAAT3 in neurons, we performed subcellular fractionation experiments in murine brain lysates to purify the light membrane fraction (containing small vesicles, e.g. endosomes), the synaptosomal fraction, the extra-synaptic membranes (plasma membrane, ER and Golgi) and the PSD1 fraction (containing the post-synaptic compartment) (Fig. S2). Similar to what had been reported for rat brain fractionations before (González et al., 2007), we mainly detected EAAT3 in the light membrane fraction, the synaptosomal fraction, and the extra-synaptic membranes of the mouse brain (Fig. S2). Remarkably, SorCS2 showed the same subcellular distribution with an obvious enrichment in the extra-synaptic membranes, the synaptosomes, and the light membrane fraction. By contrast, we did not observe enrichment of SorCS2 in PSD1 characterized by the presence of GluA1, GluA2, and PSD95 (Fig. S2). In agreement with these fractionation data, immunostainings in cultured hippocampal neurons showed that EAAT3 and SorCS2 were both present in early endosomes and recycling endosomes, characterized by Rab5 and Rab11, respectively (Fig. S2).

Jointly, our proteomics data and the reported link between mGlur3 and JWA with glutamate signaling and glutathione synthesis, suggested that the glutamate/cysteine transporter EAAT3 may represent a direct target of SorCS2-dependent sorting in neurons. In support of this hypothesis, SorCS2 and EAAT3 were expressed in the same neuronal populations in the mouse brain *in vivo*. In detail, immunostaining revealed that expression patterns for SorCS2 and EAAT3 largely overlapped, in particular in the pyramidal neurons of the cortex and in neurons of CA2 and dentate gyrus (DG) of the hippocampus (Fig. 5 A). EAAT3 is expressed both in excitatory and in inhibitory GABA-ergic neurons (Conti et al., 1998). However, we did not observe SorCS2 expression in GABA-ergic, GAD67-positive neurons, neither in the mouse brain nor in neuronal cultures (Fig. S3), indicating that co-expression of EAAT3 and SorCS2 occurs in excitatory neurons. An overlap in the expression patterns of SorCS2 and EAAT3 was also visible in the human hippocampus in healthy (Fig. 5 B) and epileptic brain tissue (Fig. S4). At the subcellular level, SorCS2 and EAAT3 immunosignals strongly overlapped in cultured hippocampal neurons and rat C6 glioma cells (Fig. 5 C).

**Figure 5.**
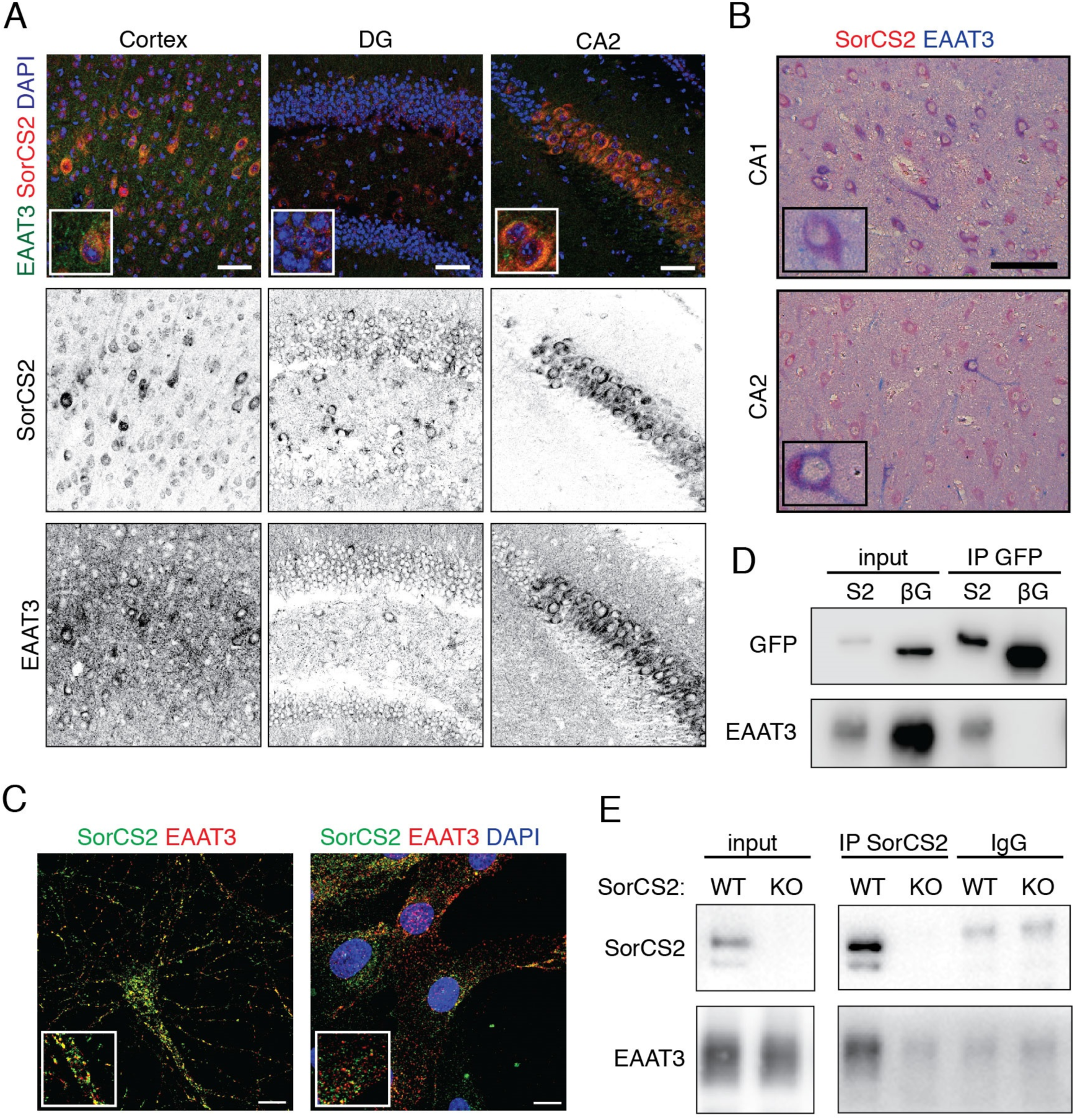
EAAT3 is a direct target of SorCS2. (A) Representative images of SorCS2 (red) and EAAT3 (green) immunostaining on mouse brain sections. Tissues were counterstained with DAPI (blue). Single confocal z-planes are shown for cortex, dentate gyrus (DG) and CA2 region. Merged images are given in the upper panel. The lower panels represent separated channels shown in greyscale. Insets show higher magnification of the given images. Scale bar: 50 μm. See also Fig. S2 and S3. (B) Immunostainings of SorCS2 (red) and EAAT3 (blue) in the healthy human hippocampus (CA1 and CA2). Insets show higher magnification of the given images. Scale bar: 100 μm. See also Figure S4. (C) Immunostaining for SorCS2 (green) and EAAT3 (red) in cultured mouse hippocampal neurons (left) and C6 cells (right). Single confocal z-planes are shown. Scale bars: 10 μm. (D) Western blot analysis of SorCS2-GFP (S2), β-galactosidase-GFP (βG), and EAAT3 in CHO cell transfected with the respective expression constructs. Expression of all three proteins is seen in cell lysates prior to immunoprecipitation (input). After IP with anti-GFP antiserum, coIP of EAAT3 with S2 was evident. No coIP of EAAT3 was seen with βG. This experiment was replicated three times. (E) Western blot analysis of endogenous SorCS2 and EAAT3 in wild-type mouse hippocampal lysates prior (input) and after SorCS2 immunoprecipitation (IP). Efficient coIP of EAAT3 was seen when using anti-SorCS2 antiserum (IP SorCS2) but not with non-specific IgG (IgG). As a negative control, no coIP of EAAT3 with anti-SorCS2 antiserum was detected from SorCS2-deficient tissue (KO). This experiment was replicated three times.

### SorCS2 interacts with EAAT3 for trafficking to the plasma membrane

To further substantiate EAAT3 as a target for SorCS2-mediated sorting, we tested interaction of the two proteins using co-immunoprecipitation (coIP). To do so, Chinese hamster ovary (CHO) cells were transiently transfected with contructs encoding EAAT3 and a fusion protein of SorCS2 with green fluorescence protein (SorCS2-GFP). Formation of a complex of EAAT3 and SorCS2 was substantiated by efficient coIP of EAAT3 with SorCS2-GFP (Fig. 5 D). No coIP of EAAT3 was seen with the control protein, GFP-tagged β-galactosidase (βG). We confirmed EAAT3 and SorCS2 interaction in tissue by coIP of the endogenous proteins from mouse hippocampal lysates of WT mice using anti-SorCS2 antisera (Fig. 5 E). No IP of EAAT3 in wild-type tissue was seen using non-immune IgG. Also, no coIP of EAAT3 was detected using anti-SorCS2 antisera in S2KO hippocampal lysates (Fig. 5 E).

So far, our data documented direct interaction of SorCS2 with EAAT3 in the brain. We hypothesized that SorCS2 may act as a sorting receptor for EAAT3, enabling it to reach the cell surface where EAAT3 exerts its function in glutamate and cysteine uptake. EAAT3 and other EAAT proteins form homomultimers that constitute the active form of this transporter (Eskandari et al., 2000; Gendreau et al., 2004; Haugeto et al., 1996). In support of this notion, we detected mono- and multimeric variants of EAAT3 in total lysates from primary mouse neurons using Western blotting (Fig. 6 A). Strikingly, in the purified neuronal cell surface fraction, we mainly detected multimeric EAAT3, whereas the monomer was barely visible (Fig. 6 A). Importantly, the levels of multimeric EAAT3 in the cell surface fraction were significantly decreased in S2KO primary neurons as compared to WT neurons (Fig. 6 A-B), implicating SorCS2 in trafficking of EAAT3. We corroborated this finding in an alternative cell model of SorCS2 deficiency by transient knock down of the receptor in the C6 cell line. Acute SorCS2 depletion in C6 cells also led to a significant decrease in EAAT3 levels (Fig. 6 D-E).

**Figure 6.**
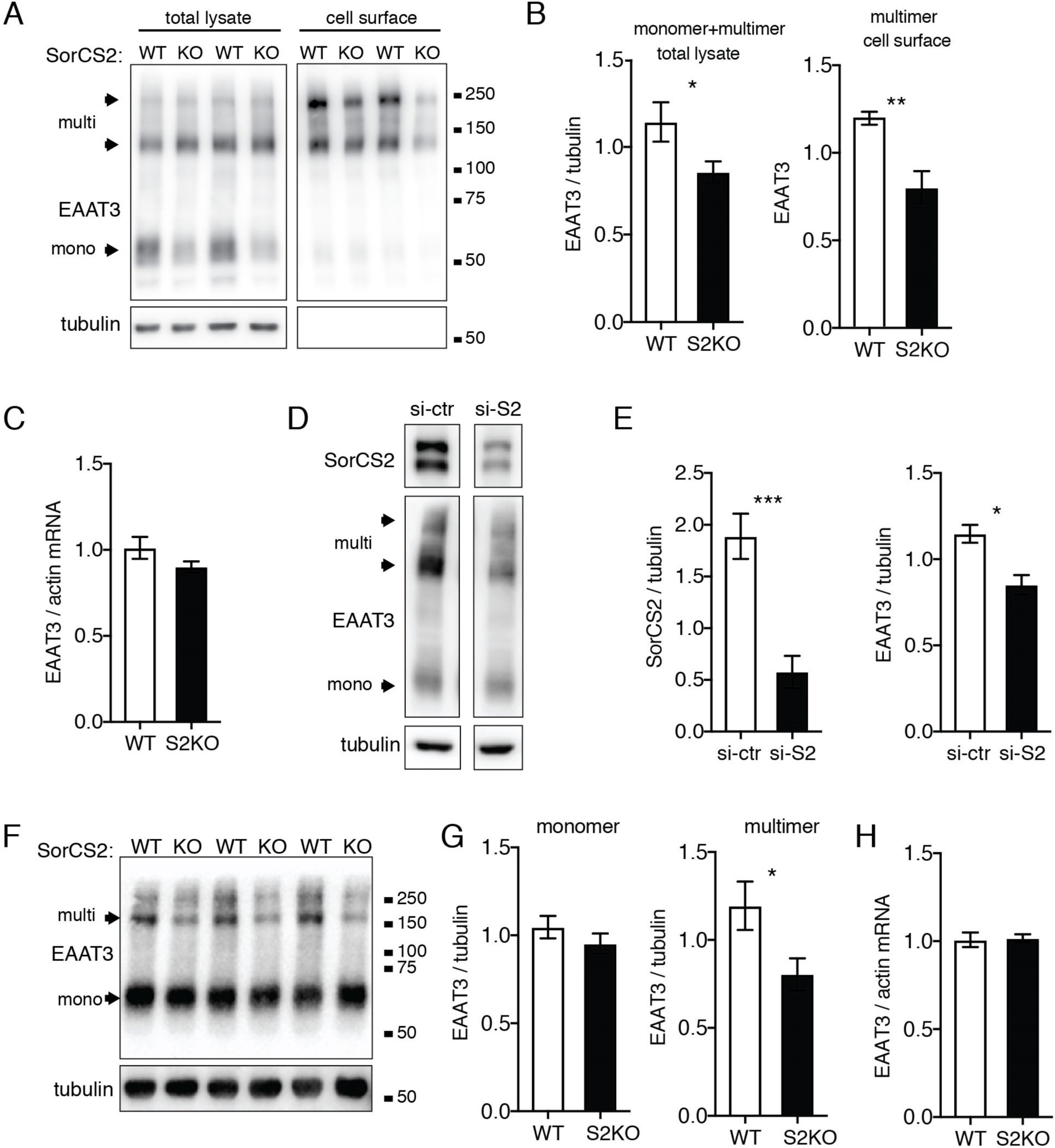
Loss of SorCS2 decreases cell surface levels of EAAT3. (A) Western blot analysis of SorCS2 and EAAT3 levels in total lysate and cell surface fractions of primary wild-type (WT) and S2KO (KO) mouse neurons. In total cell lysate, EAAT3 is visible both as monomeric and multimeric forms (arrowheads). By contrast, multimers represent the predominant form of EAAT3 present at the cell surface. Detection of tubulin served as a loading control. (B) Quantification of EAAT3 levels in WT and S2KO neurons. For total lysates, signal intensities for EAAT3 monomers and multimers were combined. For the cell surface fraction, only levels of the EAAT3 multimer were scored. EAAT3 levels in the total lysate were normalized to tubulin levels. Mean ± SEM; n=7 independent neuronal preparations per genotype; *, p<0.05; **, p<0.01 (unpaired t-test). (C) EAAT3 mRNA levels in WT and S2KO primary neurons as assessed by qRT-PCR (relative to actin mRNA). Mean ± SEM; n=5-6 independent neuronal preparations per genotype. (D) Western blot analysis of SorCS2 and EAAT3 levels in C6 cell lysates 48 hours after transfection with siRNA directed against SorCS2 (si-S2) or with a non-targeting control siRNA (si-ctr). Monomeric and multimeric forms of EAAT3 in the cell lysate are indicated by arrowheads. Detection of tubulin served as loading control. (E) Quantification of levels of SorCS2 and EAAT3 (mono- and multimeric forms) in C6 cell lysates 48 hours after transfection with control (si-ctr) or anti-SorCS2 siRNA (si-S2) (normalized to tubulin) as exemplified in panel D. Mean ± SEM; n=9 independent transfection experiments; *, p<0.05; ***, p<0.001 (unpaired t-test). (F) Representative immunoblot showing EAAT3 levels in hippocampal lysates from WT and S2KO mice (KO). Tubulin is shown as a loading control. Monomeric and multimeric forms of EAAT3 are indicated by arrowheads. (G) Quantification of levels of EAAT3 mono- and mulitmers in WT and KO mouse hippocampal lysates (normalized to tubulin levels) as exemplified in panel F. Mean ± SEM; n=10 mice per genotype; *, p<0.05 (unpaired t-test). (H) EAAT3 mRNA levels in WT and S2KO hippocampi as assessed by qRT-PCR (relative to actin mRNA). Mean ± SEM; n=9 mice per genotype.

Finally, we confirmed changes in subcellular distribution of EAAT3 in the mouse hippocampus *in vivo*. In these experiments, we assumed that the multimeric form of EAAT3 in the hippocampal lysates represented the cell surface-exposed transporter, as shown for primary neurons before (Fig. 6 A). In line with our findings in cultured neurons and C6 cells, levels of the multimeric form of EAAT3 were decreased in hippocampal extracts from S2KO mice, whereas the levels of the EAAT3 monomer remained unchanged (Fig. 6 F-G). Loss of SorCS2 did not impact the levels of *Eaat3* transcript, neither in cultured neurons (Fig. 6 C) nor in brain tissue (Fig. 6 H), indicating that the observed alterations in EAAT3 cell surface exposure were the consequence of perturbation of a post-transcriptional mechanism, probably impaired sorting to the cell surface in SorCS2-deficient neurons.

### SorCS2 deficiency leads to EAAT3 missorting

The extent of cell surface exposure of EAAT3 is determined by cell surface delivery, endocytosis, and recycling through endosomal compartments (Yang and Kilberg, 2002). In our studies described above, loss of SorCS2 resulted in decreased cell surface exposure of EAAT3, but the fate of missorted EAAT3 molecules in SorCS2-deficient neurons remained elusive. Conceptually, once unable to reach its proper destination at the plasma membrane, EAAT3 may be redirected to another subcellular compartment. In line with this assumption, we observed that the fraction of EAAT3 colocalizing with the late endosome marker Rab7 was significantly larger in cultured S2KO neurons as compared to WT cells (Fig. 7 A-B). To test whether missorting of EAAT3 to late endosomes in SorCS2 mutant neurons resulted in aberrant lysosomal catabolism of the transporter, we quantified the amount of EAAT3 in the cell surface fraction of cells treated with a lysosomal inhibitor cocktail (leupeptin, pepstatin, and chloroquine). Blocking lysosomal proteolysis caused a significant increase in cell surface levels of multimeric EAAT3 in S2KO neurons (Fig. 7 C-D). This finding clearly pointed to a state of enhanced proteolytic degradation of EAAT3 in SorCS2-deficient neurons.

**Figure 7.**
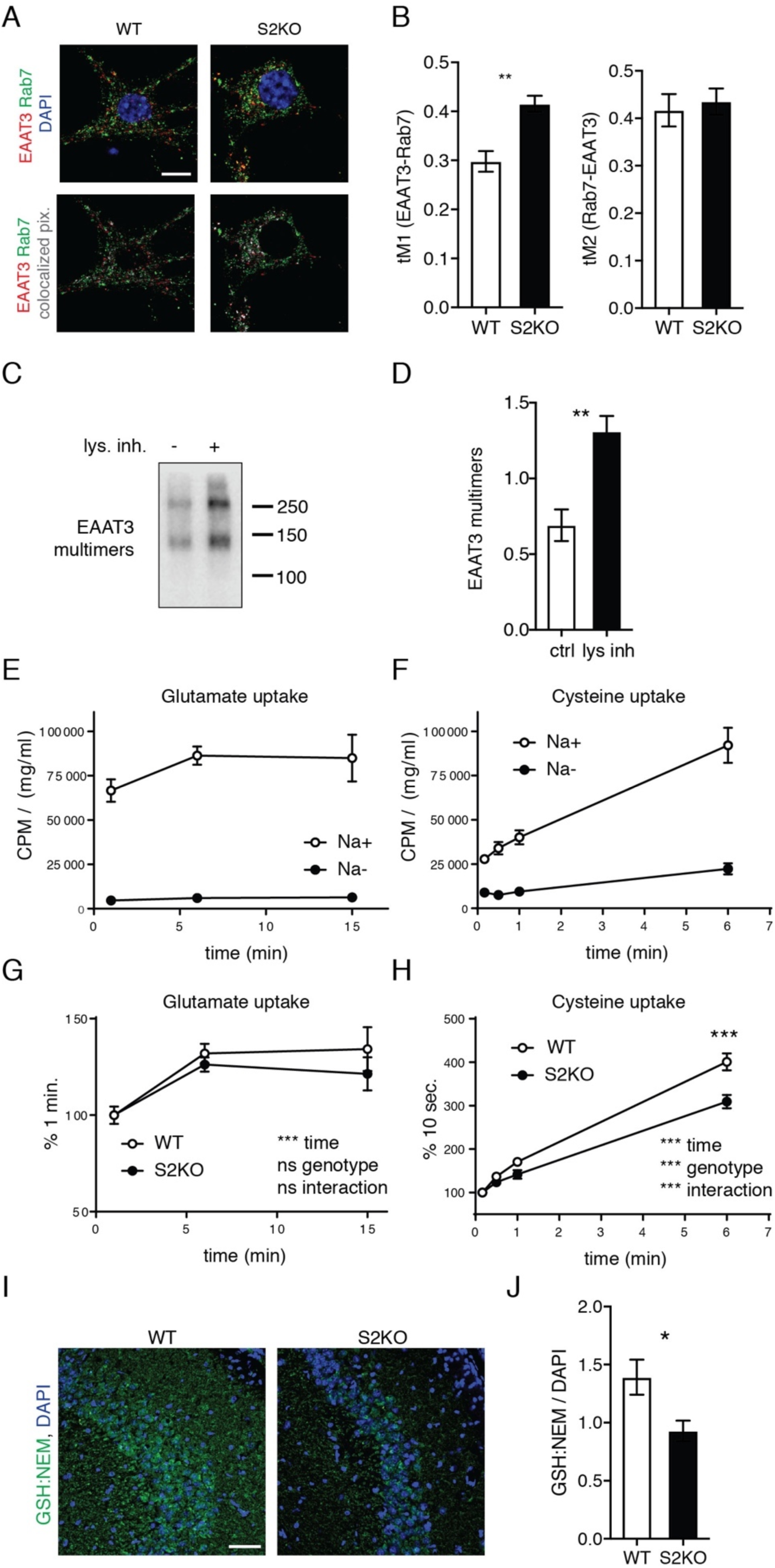
SorCS2 deficiency leads to EAAT3 mis-sorting and disrupts cysteine import into neurons. (A) Representative confocal images showing immunostaining of EAAT3 (red) and late endosome marker Rab7 (green) in primary WT and S2KO hippocampal neurons. Cells were counterstained with DAPI. Merged images of the immunosignals are given in the upper panels. In the lower panels, colocalizing EAAT3 and Rab7-positive pixels are depicted in grey. Single confocal z-planes are shown. Scale bar: 10 μm. (B) Quantification of the extent of colocalization of EAAT3 and Rab7 in WT and S2KO neurons (Manders coefficients, tM1: fraction of EAAT3 signal overlapping with Rab7 signal; tM2: fraction of Rab7 signal overlapping with EAAT3). In S2KO neurons the fraction of EAAT3 present in the Rab7 compartment (tM1) is larger than in WT cells. n=10 neurons per genotype; **, *p*>0.01 (unpaired t-test). This experiment was replicated 3 times. (C) Representative Western blot analysis of EAAT3 levels in the cell surface fraction of S2KO neurons under control conditions (lys. inh.−) or after 2 hours of treatment with lysosomal inhibitors (lys. inh.+). (D) Quantification of cell surface levels of EAAT3 in S2KO cultured neurons under control conditions (ctrl) or after 2 hours of treatment with lysosomal inhibitors (lys. inh.+). Mean ± SEM, n=5 independent neuronal preparations; **, *p*<0.01 (unpaired t-test). (E) [3H]-Glutamate uptake into primary wild-type neurons in Na-containing (Na+) and Na− deprived (Na−) buffer, normalized to protein content in the neuronal lysates. (F) [35S]-Cysteine uptake into primary wild-type neurons in Na-containing (Na+) and Na− deprived (Na−) buffer, normalized to protein content in the neuronal lysates. (G) [3H]-Glutamate uptake into primary WT and S2KO neurons, normalized to protein content in the neuronal lysates and shown as percent of the value at 1 minute. Mean ± SEM, n=7-8 independent neuronal preparations per genotype; ***, -p<0.001 (two-way ANOVA). (H) Na-dependent [35S]-Cysteine uptake into primary WT and S2KO neurons, normalized to protein content in the neuronal lysates and shown as percent of the value at 10 seconds. Na− independent uptake was subtracted from the results prior to calculating final values (% at 10 sec.). Mean ± SEM, n=9 independent neuronal preparations per genotype; ***, p<0.001 (two-way ANOVA with Sidak’s multiple comparisons test). (I) Representative images showing immunostaining for glutathione:NEM adduct (GSH:NEM) in the CA2 regions of WT and S2KO mice. Scale bar: 50 μm. (J) Quantification of the GSH:NEM signal intensity in the CA2 region normalized to DAPI intensity. Mean ± SEM, n=5-6 mice per genotype; *, *p*<0.05 (unpaired t-test).

### SorCS2 is required for cysteine import into neurons

EAAT3 is a sodium-dependent transporter for glutamate and cysteine. To evaluate the impact of EAAT3 missorting upon loss of SorCS2 on ligand uptake, we measured glutamate and cysteine uptake in primary neurons from wild-type and SorCS2-deficient mice. [S35]-labeled cysteine or [H3]-labeled glutamate were applied to cultured WT and S2KO neurons and their intracellular content was assayed over time by scintillation counting. As EAAT3 is a sodium-dependent transporter, we performed parallel assays in Na-containing (Na+) and Na-deprived (Na-) buffer to control for the contribution of Na-independent transport to glutamate and cysteine uptake. No substantial contribution of Na-independent mechanisms to glutamate import into cultured neurons was noted (7% of total glutamate uptake at 6 minutes; Fig. 7 E). For cysteine, Na-independent transport accounted for 24% of total uptake at 6 minutes (Fig. 7 F). Therefore, in all subsequent experiments we corrected for its contribution to overall cysteine import to quantify exclusively the Na-dependent transport activity. Comparing WT and S2KO neurons, glutamate uptake did not show any difference between genotypes (Fig. 7 G), likely explained by the fact that cultured neurons express EAAT3 but also other Na-dependent glutamate transporters (Chen and Swanson, 2003). Contrasting glutamate uptake, Na-dependent cysteine import was significantly decreased in S2KO cultures (Fig. 7 H), documenting a functional consequence of EAAT3 missorting in SorCS2-deficient neurons. Impaired EAAT3-dependent cysteine import into neurons results in reduced glutathione synthesis in this cell type (Aoyama et al., 2006). In line with this concept, glutathione levels in the CA2 region of SorCS2 deficient mice were significantly reduced as compared to WT mice (Fig. 7 I-J), as shown by immunodetection of the glutathione-*N*-ethylmaleimide adducts (GSH:NEM; Escartin et al., 2011; Miller et al., 2009).

In conclusion, we have identified a unique role for SorCS2 in targeting of EAAT3 to the neuronal cell surface. Loss of SorCS2 impaired EAAT3-dependent uptake of cysteine and reduced glutathione levels, compromising neuronal protection against epilepsy-induced oxidative stress.

## DISCUSSION

Amino acid transporter EAAT3 is the main cysteine uptake route in neurons acting as a rate-limiting step for synthesis of the antioxidant glutathione. Although EAAT3 was initially described as a glutamate transporter, emerging evidence argues that its major function is cysteine import (Aoyama et al., 2006), as glutamate clearance *in vivo* is mainly exerted by glia. In line witht this conclusion, EAAT3-deficient mice show glutathione deficiency and age-related neurodegeneration (Aoyama et al., 2006). We now document a role for the VPS10P domain receptor SorCS2 in controlling intracellular trafficking and activity of EAAT3. Our data suggest that SorCS2 facilitates EAAT3 sorting to the cell surface to promote cysteine import for glutathione synthesis, and to protect neurons from oxidative stress, as in epilepsy.

Intracellular trafficking of EAAT3 has been studied extensively, identifying an exocytic and a recycling route taken by this transporter. The exocytic route is used by EAAT3 during biosynthesis and includes passage through ER and Golgi to the plasma membrane. It also entails glycosylation and assembly of EAAT3 into multimers (Farhan et al., 2006; Yang and Kilberg, 2002). On the other hand, the recycling route is crucial for dynamic regulation of cell surface levels of EAAT3 and involves clathrin-dependent endocytosis followed by recycling through the Rab11+ compartment back to the plasma membrane (González et al., 2007).

Proteins known to influence exocytic trafficking of EAAT3 are Reticulon2B and JWA, which regulate ER exit of EAAT3 (Liu et al., 2008; Ruggiero et al., 2008). Although JWA levels are increased in the mouse brain as a secondary consequence of SorCS2-deficiency (Fig. 4), it is unlikely that SorCS2 partakes in exocytic trafficking of EAAT3 at the ER. VPS10P domain receptors are produced as inactive pro-receptors that require proteolytic processing of a pro-peptide in the Golgi for activation (Glerup et al., 2014; Munck Petersen et al., 1999; Quistgaard et al., 2009). Thus, these receptors typically act in post-Golgi trafficking processes.

Rather than being involved in exocytosis, SorCS2 seems to be required for recycling of EAAT3. Proteins involved recycling of EAAT3 identified so far belong to the core components of intracellular membrane sorting and fusion machineries, including SNARE proteins SNAP23 (Fournier and Robinson, 2006) and syntaxin 1a (Yu et al., 2006), small GTPase Rab11 (González et al., 2007), clathrin adaptor Numb (Su et al., 2016), scaffold protein PDZK1, and clathrin adaptor complex AP2 (D’ Amico et al., 2010). Of note, Rab11 dysfunction slowed EAAT3 trafficking to the cell surface and led to impairments in cysteine uptake and glutathione synthesis (Li et al., 2010), pointing to Rab11+ endosomes as a compartment crucial for EAAT3 activity. In agreement with previous studies, we showed that a considerable fraction of EAAT3 resides in Rab11+ recycling endosome (Fig. S2). Now, we uncovered a mechanism of receptor-mediated sorting of EAAT3 crucial for its targeting to the plasma membrane. Our data suggest a model in which EAAT3 molecules internalized from the cell surface interact with SorCS2 in Rab11+ compartment to facilitate recycling to the cell surface. In SorCS2-deficient neurons, this recycling route is diminished and internalized EAAT3 molecules are redirected to Rab7+ late endosome, followed by proteolytic degradation in lysosomes. Lysosomal degradation of EAAT3 is a physiological mechanism to regulate turnover of EAAT3 (Yang and Kilberg, 2002). However, this trafficking fate seems to be overrepresented in neurons lacking the sorting receptor SorCS2.

In neurons, glutamate and cysteine transport capacity is dynamically regulated by adjusting EAAT3 levels at the cell surface. For example, plasma membrane exposure of EAAT3 decreases after kainic acid stimulation (Furuta et al., 2003; Yu et al., 2006) but increases upon activation of the PDGF pathway, PI3K, or PKC (Davis et al., 1998; Fournier et al., 2004; Guillet et al., 2005), the latter being presumably the most potent modulator of EAAT3 trafficking. Whether PKC or other signal transducers promote cell surface sorting of EAAT3 by increasing the level or acitivity of SorCS2 is unclear, but PKC activation was shown to increase phosphorylation of the carboxyl terminal domain of SORLA (Lane et al., 2010) and to induce cell surface expression of sortilin (Navarro et al., 2002). As SorCS2 shares structural similarity with these related VPS10P domain receptors, it may also be regulated by PKC to modulate EAAT3 recycling under stress conditions, such as oxidative stress known to activate PKC (Gopalakrishna and Jaken, 2000).

Our data suggest that SorCS2 is crucial for protection against oxidative damage and cell death in the PTZ kindling model of epilepsy. Oxidative stress is a hallmark of epilepsy-driven pathology and represents one of the major processes underlying the damaging effects of epileptic activity in the brain (Martinc et al., 2014). Increased levels of oxidative stress have been observed in virtually all epilepsy models (Shin et al., 2011) and in epileptic human brains (López et al., 2007; Rumià et al., 2013). Of note, increased oxidative damage and cell death in SorCS2 deficient mice coincided with their accelerated mortality. The exact link between oxidative stress, cell death, and seizure-related mortality in the S2KO mouse remains to be established. However, there is experimental evidence for a causative role of oxidative brain damage in mortality as treatment with antioxidants significantly reduced oxidative stress and improved survival in models of epilepsy (Pauletti et al., 2017; Pearson-Smith et al., 2017). Likely, epilepsy-induced mortality in S2KO mice results from oxidative damage to multiple brain regions. Although we mainly focused on the hippocampus, our studies do not exclude a role for SorCS2 in protection against oxidative stress in other areas of the brain, such as the piriform cortex or the amygdala where the receptor is also expressed (Hermey et al., 2004). Also, the role of SorCS2 in protection from oxidative stress may not be restricted to epilepsy but might be relevant for multiple brain pathologies, possibly explaining the association of this receptor with psychiatric and neurodegenerative diseases (Alemany et al., 2015; Baum et al., 2008; Christoforou et al., 2011; Ollila et al., 2009; Reitz et al., 2013). For example, upregulation of brain EAAT3 was reported not only in animal models of epilepsy and in the human epileptic hippocampus (Crino et al., 2002; Ross et al., 2011), but also in patients suffering from schizophrenia (Bauer et al., 2008). Futhermore, aberrant localization of EAAT3 was noted in the brain of Alzheimer’s disease patients, particularly in CA2 (Duerson et al., 2009). Finally, in mouse models of Huntington’s disease impaired SorCS2 function was observed (Ma et al., 2017), as well as aberrant EAAT3 localization leading to increased levels of oxidative stress (Li et al., 2010).

In summary, we identified a novel molecular mechanism of SorCS2 action in neurons that is of major relevance for brain pathophysiology. This newly discovered role in sorting of EAAT3 complements other neuronal activities of this multifunctional receptor, such as sorting of NMDA receptors (Ma et al., 2017) or regulation of signaling by the proforms of brain derived neurotrophic factor (Anastasia et al., 2013; Glerup et al., 2014, 2016) and nerve growth factor (Deinhardt et al., 2011). Likely, impairement of SorCS2 activity results in alterations of multiple neuronal sorting processes, collectively contributing to various brain pathologies associated with this receptor.

## EXPERIMENTAL PROCEDURES

### Human subjects

The cases included in this study were obtained from the archives of the Departments of Neuropathology of the Academic Medical Center (AMC) Amsterdam, The Netherlands. Hippocampal specimens from 6 patients undergoing surgery for drug-resistant TLE were examined. The tissue was obtained and used in accordance with the Declaration of Helsinki and the AMC Research Code provided by the Medical Ethics Committee. All cases were reviewed independently by two neuropathologists and the classification of hippocampal sclerosis was based on analysis of microscopic examination as described by the International League Against Epilepsy (Blümcke et al., 2013). Hippocampal control tissues were obtained during autopsy of age-matched individuals without a history of seizures or other neurological diseases (n=6). Table S1 summarizes the clinical characteristics of patients and controls.

### Mouse model

Mice with targeted disruption of *Sorcs2* (S2KO) were described before (Glerup et al., 2014). S2KO mice on an inbred C57BL/6N background were used. For *in vivo* studies, mouse strain was kept by breeding of heterozygous animals and the WT and S2KO littermates were used in experiments. Newborn mice for preparing neuronal cultures we obtained by breeding of homozygous WT (C57BL/6N) or S2KO animals.

PTZ kindling experiments were performed using male mice at the age of 8-12 weeks. In all the other studies, brain tissue was collected from the mice of both sexes at the age of 8-12 weeks.

All animal experimentation was performed in accordance with institutional guidelines following approval by the local authorities of the State of Berlin (X9012/12, O0120/13) and by the First Ethical Committee in Warsaw (209/2016).

### Neuronal cultures

Primary hippocampal or mixed cortical/hippocampal (referred to as ‘cortical’) neuronal cultures were prepared from newborn S2KO and WT mice (postnatal day 0-1) using enzymatic digestion with papain. Cortical neurons were plated on plastic plates coated with poly-D-lysine (Corning BIOCOAT) at a density of 64,000 cells/cm^2^ (biotinylation studies) or 83,000 cells/cm^2^ (neuronal stimulation). For immunocytochemistry, hippocampal neurons were plated on glass coverslips coated with poly-D-lysine and laminin at a density of 25,000 cells/cm^2^. Neurons were maintained in Neurobasal medium (Invitrogen) supplemented with B27, GlutaMAX, and penicillin/streptomycin (Invitrogen).

### Cell lines

The C6 rat glioma cell line was purchased from Culture Collections (UK). The cells were maintained in DMEM medium supplemented with 5% FBS and penicillin/streptomycin (Invitrogen). CHO cells were obtained from ATCC and grown in DMEM medium supplemented with 10% FBS and penicillin/streptomycin (Invitrogen).

### Plasmids, oligonucleotides, and cell transfections

Plasmids encoding rat EAAT3 and GFP-tagged β-Galactosidase were gifts from Michael Robinson and Jacek Jaworski, respectively (González et al., 2002; Swiech et al., 2011). The plasmid encoding murine SorCS2 tagged with GFP at the C-terminus was generated in-house. Lipofectamine ^TM^ 2000 was used for transfection of cells.

For transient knockdown of SorCS2 expression, Dharmacon siRNA (ON-TARGET plus SMART pool, L-085157-02-0005) or non-targeting control siRNA were used (ON-TARGET plus non-targeting Pool D-001810-10-05). Cells were transfected with siRNAs using X-tremeGENE siRNA transfection reagent (Roche).

### Neuronal treatments

Neuronal cultures at day *in vitro* (DIV) 10-12 were used. For NMDA treatment, the culture medium was replaced with fresh Neurobasal medium supplemented with B27 and 20 μM NMDA (Sigma-Aldrich) and the cells were incubated for 5 minutes at 37°C. In case of bicuculline treatment, bicuculline (Abcam) was added to the conditioned cell culture medium at a final concentration of 20 μM and cells were incubated for 30 minutes at 37°C. Control cells were left untreated. After the stimulation, cell culture plates were placed on ice and washed 2 times with ice-cold TBS-Ca-Mg buffer (20 mM Tris pH 7.6, 150 mM NaCl, 0.5 mM MgCl2, 1 mM CaCl_2_). Next, neurons were scraped in lysis buffer containing 20 mM Tris pH 7.5, 150 mM NaCl, 2 mM EDTA, 0.5% Triton-X, 0.5% NP40, 2 mM MgCl_2_, 10% glycerol, and protease and phosphatase inhibitors. The lysate was centrifuged (16,100x g, 15 min) to remove cell debris. The protein content was measured by BCA method and samples were prepared for Western blotting.

To block lysosome function, the culture medium was supplemented with pepstatin (10 μM), leupeptin (100 μM), and chloroquine (50 μM), and the cells were incubated for 2 hours at 37°C prior to cell surface biotinylation.

### Purification of neuronal cell surface proteins

Cortical neuronal cultures prepared from newborn WT or S2KO mice were plated on poly-D-lysine-coated plates and cell surface biotinylation was performed at DIV10-12. To do so, cells were cooled down on ice for 15 min and washed 3 times with PBS-Mg-Ca (PBS supplemented with 0.5 mM MgCl_2_ and 1 mM CaCl_2_). Next, cells were incubated on ice with EZ-Link™ Sulfo-NHS-SS-Biotin solution (Thermo Scientific; 0.5 mg/ml in PBS) for 25 min, washed 3 times with quenching buffer (40 mM glycine, 0.4% BSA in TBS-Ca-Mg) and 2 times with TBS-Ca-Mg. Cells were scraped in lysis buffer (50 mM Tris pH 7.5, 1 mM EDTA, 2 mM EGTA, 150 mM NaCl, 1% NP40, 0.5% DOC, 0.1% SDS, protease and phosphatase inhibitors) and the lysates were rotated for 1.5 hours at 4°C. After centrifugation (15 min, 14,000x *g*), the supernatants were used for pull down of biotinylated proteins with NeutrAvidin slurry (Thermo Scientific). After washing with lysis buffer, the beads were snap-frozen and stored at −80°C until mass spectrometry analysis or boiled in Laemmli buffer to release captured proteins for Western blot analysis.

### Mass spectrometry analysis of neuronal cell surface proteins

Mass spectrometry analysis was performed as described (Subkhangulova et al., 2018). Samples containing biotinylated cell surface proteins were run on a stacking SDS-PAGE collecting all proteins in a single band. After Coomassie staining, the gel pieces were minced and digested with trypsin using a PAL robot (Axel Semrau/CTC Analytics). Peptides were extracted with extraction buffer (80% acetonitrile, 0.1% [v/v] formic acid) and dried in a speed-vac, followed by purification on C18 stage-tips. The eluted peptides were dried in a speed-vac and resuspended in 3% acetonitrile, 0.1% (v/v) formic acid.

The samples were measured by LC-MS/MS on a Q-Exactive Plus mass spectrometer (Thermo) connected to a Proxeon easy-nLC system (Thermo). The peptides were separated on an in-house prepared nano-LC column (0.074 mm × 250 mm, 3 μm Reprosil C_18_, Dr Maisch GmbH) using a flow rate of 0.25 μl/min. MS acquisition was performed at a resolution of 70,000 in the scan range from 300 to 1,700 m/z. MS2 scans were carried out at a resolution of 15,500 with the isolation window of 2.0 m/z. Dynamic exclusion was set to 30 s, and the normalized collision energy was specified to 26 eV.

For analysis, the MaxQuant software package version 1.5.2.8 was used. A FDR of 0.01 was applied for peptides and proteins, and the Andromeda search was performed using a *Mus musculus* Uniprot database (August 2014). MS intensities were normalized by the MaxLFQ algorithm implemented in MaxQuant. MaxLFQ-normalized intensities among the replicates of the groups to be related were used for statistical comparison. Proteins were considered as specifically enriched for a group if they fulfilled a defined fold change (FC) of the averaged normalized intensities (threshold value ±0.5 for log2(S2KO/WT) and a *P*-value from a Student’s *t*-test < 0.05 for comparison of two groups (threshold value 1.3 for −log_10_(*P*-value)). For data visualization, the −log_10_(*P*-value) was plotted against the log2(FC) using *R* (http://www.r-project.org). For proteins detected in either S2KO or WT samples only, the log_2_(S2KO/WT) was set to 10 or −10, respectively.

### PTZ kindling procedure

For up to 5 weeks, mice were intraperitoneally injected with subconvulsive doses (40 mg/kg body weight) of pentylenetetrazol (PTZ) in saline between 9:00 and 11:00 a.m. three times a week as described (Becker et al., 1992). After each injection, the convulsive behavior was video recorded for 30 min and the resultant seizures were scored as follows: stage 0, no response; stage 1, ear and facial twitching; stage 2, convulsive waves axially through the body; stage 3, myoclonic jerks and rearing; stage 4, turning over onto the lateral position; stage 5, turning over into the back and generalized tonic-clonic seizures and 6, for animals that died during the seizure. The animals were considered to be kindled after having had at least three consecutive sessions when they reached stage 5 seizures. After completion of the experiment (15 sessions) or when animals died during the seizure, their brains were snap frozen on dry ice.

### Immunohistochemistry

Human brain tissue was fixed in 10% buffered formalin and embedded in paraffin. Paraffin-embedded tissue was sectioned at 5 μm, mounted on pre-coated glass slides (Star Frost, Waldemar Knittel) and processed for immunohistochemical staining. Sections were deparaffinated in xylene, rinsed in ethanol (100%, 95%, 70%), and incubated for 20 minutes in 0.3% hydrogen peroxide diluted in methanol. Antigen retrieval was performed using a pressure cooker in 10 mM sodium citrate, pH 6.0 at 120°C for 10 minutes. Slides were washed with phosphate-buffered saline (PBS; 0.1 M, pH 7.4) and incubated overnight with primary antibody (rabbit anti-SorCS2, LS-C501334, Life Span Biosciences 1:450) in Normal Antibody Diluent (Immunologic, Duiven, The Netherlands) at 4°C. After washing in PBS, sections were stained with a polymer based peroxidase immunohistochemistry detection kit (Brightvision plus kit, ImmunoLogic) according to the manufacturer’s instructions. Staining was performed using Bright DAB substrate solution (ImmunoLogic, Duiven, the Netherlands). Sections were dehydrated in alcohol and xylene, and coverslipped.

Double-labeling of SorCS2 with EAAT3 (rabbit, Cell Signaling #14501, 1:150) was performed as previously described (Iyer et al., 2013). Sections were incubated with Brightvision poly-alkaline phosphatase (AP)-anti-rabbit (Immunologic) for 30 minutes at room temperature, and then washed with PBS. AP activity was visualized with the AP substrate kit III Vector Blue (SK-5300, Vector laboratories Inc.). To remove the first primary antibody, sections were incubated at 121°C in citrate buffer (10 mM NaCi, pH 6,0) for 10 min. Incubation with the second primary antibody was performed overnight at 4°C. AP activity was visualized with the alkaline phosphatase substrate kit I Vector Red (SK-5100, Vector laboratories Inc.). Negative control sections incubated without the primary antibodies or with the primary antibodies, followed by heating treatment were essentially blank.

Mouse brains were dissected from 8 to 12 weeks old mice intracardially perfused with phosphate-buffered saline (PBS) and 4% paraformaldehyde (PFA) in PBS. After post-fixation (PFA, overnight) and cryopreservation in 30% sucrose/PBS, brains were cut in 50 μm coronal sections using a sliding microtome. Free-floating sections were blocked in 1% horse serum in PBS and incubated with primary followed by secondary antibodies diluted in PBS, supplemented with 1% bovine serum albumin (BSA), 1% normal donkey serum (NDS) and 0.5% Triton-X.

After PTZ kindling experiments, mouse brains were dissected and rapidly frozen. Next, 25 μm coronal sections were cut using a cryostat, immediately mounted on glass slides, and kept at −20°C until further use. Prior to staining, the sections were thawed at room temperature for 30 min and fixed in 4% PFA/PBS for 30 min. Sections were blocked and incubated with antibodies as described for free-floating sections above. For detection of 8OHdG, the blocking buffer contained 10% NDS, 1% BSA, and 0.3% Triton-X in TBS-T (132 mM NaCl, 2.7 mM KCl, 25 mM Tris pH 7.5, 0.1% Tween-20) and antibodies were diluted in TBS-T.

TUNEL reaction was performed using a commercially available kit (Roche). Sections were thawed for 30 min at 4°C and fixed in 25% acetic acid in ethanol for 30 min at 4°C. Next, the sections were blocked with 2% BSA, 1.5% normal goat serum, 0.1% Triton-X in PBS, and subjected to TUNEL reaction as described in the manufacturer’s manual.

To visualize glutathione, the sections were first incubated for 4 hours at 4°C in 10 mM N-ethylmaleimide (NEM)/PBS. After washing with PBS, the sections were incubated overnight at 4°C with anti-GSH:NEM antibody in PBS supplemented with 0.3% Trition-X. Next, they were washed and incubated for 2 hours at room temperature with fluorescently labeled secondary antibody.

C6 cells grown on glass coverslips were fixed with 4% PFA/PBS. Hippocampal neurons grown on glass coverslips coated with PDL and laminin were fixed with 4% PFA/PBS at DIV 10-12. Next, the cells were washed with PBS, blocked for 1 hour in PBS supplemented with 5% NDS and 0.3% Triton-X, and incubated with primary antibodies diluted in PBS with 1% BSA and 0.3% Triton-X.

Following antibodies were used for immunostainings: anti-EAAT3 (Cell Signaling, CS14501, 1:100; Santa Cruz Biotechnology, SC7761, 1:100), anti-GAD67 (EMD Millipore, MAB5406, 1:100), anti-Rab5 (Cell Signaling, CS2143, 1:100), anti-Rab7 (Cell Signaling, CS9367, 1:100), anti-Rab11 (Cell Signaling, CS-D4F5, 1:100), anti-SorCS2 (R&D, AF4237, 1:100), anti-GSH:NEM (Millipore, clone 8.1GSH, 1:100). Primary antibodies were visualized using Alexa Fluor 555, 488 or 647 secondary antibody conjugates. 8OHdG was immunodetected using FITC-conjugated antibody (Santa Cruz Biotechnology, SC93871, 1:50) and GFAP was detected using Cy-3 conjugated antibody (Sigma-Aldrich C9205, 1:1,500). The cells and tissue sections were counterstained with DAPI and mounted with DAKO fluorescence mounting medium.

### Hippocampal lysates

Mouse hippocampi were homogenized in buffer containing 20 mM Tris pH 7.5, 150 mM NaCl, 1 mM MgCl_2_, 1 mM CaCl_2_, and protease and phosphatase inhibitors. The lysates were kept for 20 minutes on ice and then centrifuged (1,000x *g*, 10 min) to remove tissue debris. Next, the supernatant was supplemented with CHAPS to a final concentration of 0.6% and rotated for 1h at 4°C. The lysates were centrifuged (16,100x *g*, 15 min) and supernatants used for Western blot analysis.

### Tissue fractionation

Brain subcellular fractionations were performed as illustrated on Figure S2. Brain tissues (hippocampi and cortex) from 8 to 12 weeks old mice were homogenized in Tris-buffered sucrose with Ca and Mg (0.32 M sucrose, 6 mM Tris, 1 mM MgCl_2_, 0.5 mM CaCl_2_, protease and phosphatase inhibitors, pH 8). The resulting suspension was centrifuged (1,400x *g*, 20 min) yielding the supernatant (S1a) and the pellet, that was homogenised again in Tris-buffered sucrose with Ca and Mg and centrifuged to remove nuclear debris (710x *g* 10 min) and to collect the supernatant (S1b). Supernatants S1a and S1b were combined to get a total lysate (S1) that was further centrifuged (13,800× *g*, 30 min) to pellet the crude membrane fraction (P2). The resulting supernatant (S2) was subjected to ultracentrifugation (100,000× *g*, 1 h) to yield the light membrane fraction (P3). The P2 fraction was homogenized in Tris-buffered sucrose (0.32 M sucrose, 6 mM Tris pH 8) and layered onto a discontinuous sucrose density gradient (0.85 M, 1.0 M, 1.2 M sucrose buffered with 6 mM Tris pH 8) and centrifuged (82,500x *g*, 2h) to recover the extra-synaptic membranes at the interphase between the layered P2 fraction and 0.85 M sucrose, and the synaptosomal fraction at the interphase between 1.0 M and 1.2 M sucrose. The synaptosomal fraction was further supplemented with Triton-X100 to a final concentration of 0.5%, and incubated on ice for 15 min. Further centrifugation (32,000x *g*, 20 min) pelleted the PSD1 fraction, that was resuspended in 40 mM Tris pH 8. Protein concentrations were measured for all the fractions and equal amounts of protein were loaded on an SDS-PAGE gel for Western blot analysis.

### Immunoprecipitations

Hippocampi from WT and S2KO mice were homogenized in lysis buffer containing 20 mM Tris pH 7.5, 150 mM NaCl, 1 mM CaCl_2_, 1 mM MgCl_2_, protease and phosphatase inhibitors, and kept on ice for 20 min. Next, the lysates were centrifuged (1,000x *g*, 10 min) and the tissue debris was discarded. CHAPS was added to the lysates to a final concentration of 0.6% and the lysates were rotated for 1 hour at 4°C. After centrifugation (16,100x *g*, 15 min), the supernatant was used for immunoprecipitation with protein G agarose beads coupled to anti-SorCS2 antibody (R&D, AF4237) or to unspecific sheep IgG.

Chinese hamster ovary cells were transfected with expression constructs encoding rat EAAT3 and GFP-tagged β-Galactosidase or murine SorCS2 using Lipofectamine™ 2000 reagent (Invitrogen). Two days after transfection, cells were washed twice and scraped in TBS-Ca-Mg buffer, and centrifuged to obtain a cell pellet (16,100x *g*, 15 min). The cell pellet was resuspended in lysis buffer (20 mM Tris pH 7.5, 150 mM NaCl, 1 mM MgCl_2_, 1 mM CaCl_2_) supplemented with 0.6% CHAPS, and protease and phosphatase inhibitors. After shaking for 30 minutes on ice, the lysate was centrifuged to remove the cell debris (16,100x *g*, 15 min). The supernatant was applied to GFP Trap beads (Chromotek) for GFP immunoprecipitation. The samples were rotated for 1 hour at 4°C before the resin was washed 3 times with lysis buffer supplemented with 0.2% CHAPS. Proteins bound to the resin were released by boiling in Laemmli buffer and analyzed by Western blot.

### Western blot

Tissue and cell lysates were analyzed by Western blot using the following antibodies: anti-SorCS2 (R&D, AF4237, 1:1000), anti-EAAT3 (Cell Signaling, CS14501, 1:1000), anti-p-ERK (Cell Signaling CS4370, 1:2000), anti-p-p38 (Cell Signaling, CS4511, 1:1000), anti-tubulin (EMD Millipore, CP06, 1:5000), anti-mGlur2/3 (Novus Biologicals, NB300-124, 1:1000), anti-actin (Abcam, ab8227, 1:2000), anti-JWA (Trans Genic, KR057, 1:250), anti-N-cadherin (Cell Signaling, CS14215, 1:1000), anti-GFP (MBL International, MBL598, 1:2000), anti-Rab11 (Cell Signaling, CS D4F5, 1:1000), anti-synaptophysin (Synaptic Systems, 101011, 1:5000), anti-GluA1 (EMD Millipore, MAB2263, 1:1000), anti-GluA2 (EMD Millipore, MABN71, 1:1000), anti-PSD95 (Cell Signaling, CS3409, 1:1000).

### Quantitative RT–PCR

Total RNA was extracted from tissue and cell lysates using TRIzol reagent and purified with RNeasy Mini Kit (Qiagen). Reversely transcribed cDNA from total RNA was subjected to qRT–PCR using the following Taqman Gene Expression Assays: Actb (actin, Mm02619580), Ar6ip5 (JWA, Mm00480826), mGlur3 (Mm00725298), EAAT3 (Mm00436590). Fold change in gene expression was calculated using the cycle threshold (CT) comparative method (2-ddCT) normalizing to Actb CT values.

### Cysteine and glutamate uptake assays

Cultures of cortical neurons from WT and S2KO mice were used at DIV 10-13.

Cells were washed 3 times with warm assay buffer (5 mM Tris pH 7.5, 10 mM HEPES, pH 7.27, 2.5 mM KCl, 1.2 mM CaCl_2_, 1.2 mM MgCl_2_, 1.2 mM K_2_HPO4, 10 mM glucose; 140 mM NaCl) and then incubated with the warm assay buffer containing L-[35S]-cysteine (0.005 uCi/ul, PerkinElmer), unlabeled L-cysteine (10 μM, Sigma-Aldrich) and DTT (100 μM, 1,4-Dithiothreitol, Sigma-Aldrich) or warm assay buffer containing L-[3,4-3H]-glutamic acid (100 nM, PerkinElmer) and unlabeled L-glutamic acid (10 μM, Sigma-Aldrich).

To study Na-independent cysteine uptake, the sodium chloride in the assay buffer was replaced with choline chloride (Sigma-Aldrich,) and the pH was adjusted with KOH.

Cysteine uptake assays were performed at room temperature for 10, 30, 60, and 360 seconds. Glutamate uptake was performed at 37°C for 0.5, 1, 6, and 15 minutes.

At indicated time points, cells were rapidly washed 3 times with cold Tris-Ca-Mg buffer followed by lysis. Cells were scraped in lysis buffer (50 mM Tris, pH 7.4, 140 mM NaCl, 1% Triton X-100) supplemented with protease inhibitor cocktail and incubated for 30 minutes on ice. Subsequently, the lysates were centrifuged at 13,400x *g* for 10 minutes and the supernatant was used for the further analysis.

The radioactive cysteine and glutamic acid contents were determined in a Liquid Scintillation Analyzer (Tri-Carb 2800TR, PerkinElmer) using 10 μl of the supernatant sample in 4 ml of liquid scintillation cocktail (Rotiszint eco plus, Carl Roth).

Values were normalized to protein content and, in case of cysteine uptake, the values obtained for Na-independent cysteine uptake were subtracted from the total uptake. Finally, values were given relative to the value obtained for the earliest time point of the uptake experiment (set to 100).

### Image quantification

Western blot signals were quantified using the Image Studio Lite software. Microscopy images were quantified using Fiji software. Colocalization analysis was performed with Colocalization Threshold plugin.

To quantify the number of TUNEL-positive cells, the CA2/3 region was manually selected, color channels were separated, and threshold was applied to minimize the background signal. Next, DAPI-positive regions were used to mask the green channel (TUNEL) and TUNEL-positive particles were counted automatically.

The 8OHdG signal intensity was quantified in manually selected CA and DG regions after separating the channels and applying a threshold to minimize the background signal. For each set of tissue sections stained in parallel, intensities were normalized to a mean value obtained for the WT mice.

The GSH:NEM signal intensity was quantified along with DAPI signal intensity within a manually selected region containing CA2 neurons. The GSH:NEM signal measurement was further normalized to DAPI intensity.

### Statistical analysis

For all *in vivo* experiments, an indicated number *n* is the number of mice per group used in an experiment. For neuronal culture experiments, an indicated number *n* is the number of independent neuronal preparations (biological replicates) used or, in case of colocalization studies, of individual cells quantified in a given experiment. Each mouse (or biological replicate/individual cell in cell culture experiments) represents a statistically independent experimental unit, which was treated accordingly as an independent value in the statistical analyses. Statistical analyses were performed using GraphPad Prism software. For comparison between two experimental groups, a two-tailed unpaired *t*-test was used. PTZ kindling response was assayed with repeated measures two-way ANOVA. Survival curve was analyzed using Gehan-Breslow-Wilcoxon test. For all other data with two independent variables (factors), two-way ANOVA with Sidak’s multiple comparisons test was applied. Where applicable, outlier analysis was performed using Grubb’s test. The details of statistical analysis are specified in the figure legends.

## AUTHOR CONTRIBUTIONS

Conceptualization, A.R.M, T.E.W, K.L., E.A.; Methodology, A.R.M, G.D.; Investigations, A.R.M., A.A., K.S., K.N., E.A.V., O.P., G.D.; Writing of the manuscript, A.R.M., T.E.W; Funding acquisition, T.E.W, E.A., K.L.; Resources, A.N.; Supervision, A.R.M., T.E.W., K.L., E.A.

## ACKNOWLEDGEMENTS

We are indepted to Jasper J. Anink, Andra Eisenmann, Tajana Pasternack, and Maria Schmeisser for expert technical assistance. This work was supported by grants from the European Research Council (BeyOND No. 335692, to TEW), the Helmholtz Association (AMPro, to TEW), the Berlin Institute of Health (Collaborative Research Group 11220008, to TEW), the European Union’s Seventh Framework Programme (FP7/2007-2013, grant agreement no. 602102, EPITARGET; to EA and KL), the European Union’s Horizon 2020 Research and Innovation Programme (Marie Sklodowska-Curie grant agreement no. 722053, EU-GliaPhD; to EA), and the Polish Ministry of Science and Education grant W19/7.PR/2014 (to KL).We thank Michael Robinson and Jacek Jaworski for providing plasmids encoding EAAT3 and GFP-tagged β-Galactosidase, respectively, and Tilman Breiderhoff for generating the plasmid encoding SorCS2-GFP.

## SUPPLEMENTAL INFORMATION

### SUPPLEMENTAL TABLES

**Table S1. Realted to Figure 1.** Clinical features of patients involved in the study.

| Pathology | n | Gender<br>m/f | Age | Duration of<br>epilepsy (years) | Age at<br>onset | Number of seizures<br>(per month) |
| --- | --- | --- | --- | --- | --- | --- |
| **Control** | 6 | 3/3 | 43.8 (31-60) | - | - | - |
| **TLE – HS** | 6 | 3/3 | 38.8 (25-62) | 22.8 (15-61) | 9.0 (1-27) | 12.8 (2-25) |

TLE, temporal lobe epilepsy; HS, hippocampal sclerosis. Values are given as mean (range).

**Table S2. Related to Figure 4.** Results of the mass spectrometry analysis of cell surface proteomes of WT and S2KO neurons. Excel file containing the complete results of the quantitative mass spectrometry analysis of protein composition of cell surface fractions derived from WT and S2KO primary neurons.

### SUPPLEMENTAL FIGURES

**Figure S1. Related to Figure 4.**
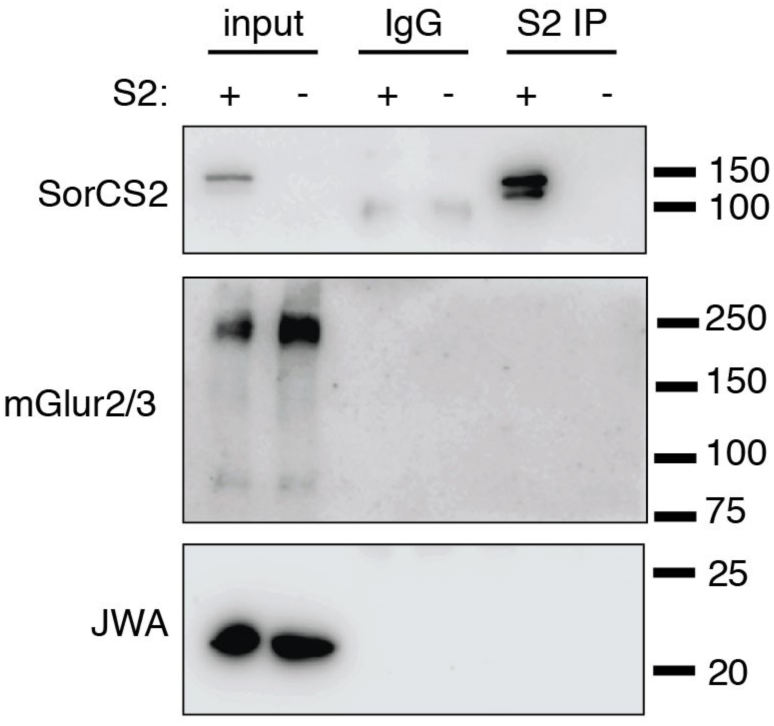
mGlur3 and JWA do not co-immunoprecipitate with SorCS2. Western blot analysis of co-immunoprecipitation experiments. SorCS2 was immunoprecipitated from mouse hippocampal lysates of wild-type (S2+) but not S2KO (S2−) mice (S2 IP). No immunoprecipation was seen in wild-type tissue using non-immune IgG (IgG). Although detected in the input sample using specific antisera, mGlur2/3 and JWA were not co-immunoprecipitated with anti-SorCS2 antibodies (S2 IP).

**Figure S2. Related to Figure 5.**
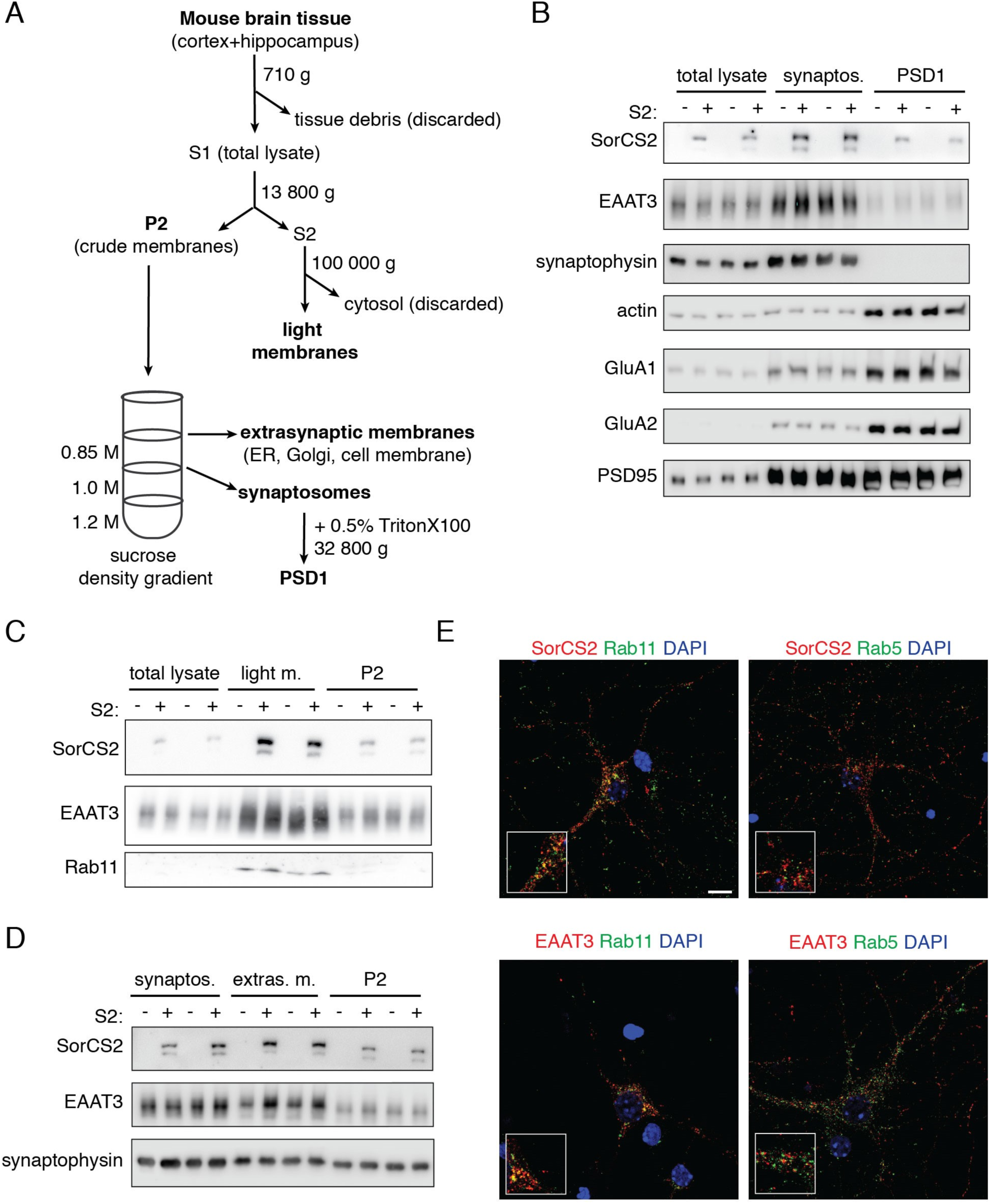
SorCS2 and EAAT3 are enriched in the same subcellular compartments in the mouse brain. (A) Schematic depiction of the subcellular fractionation procedure. Mouse brain tissue (cortex and hippocampus) was subjected to serial centrifugations to obtain P2, light membranes, extra-synaptic membranes, synaptosomes, and post-synaptic densities (PSD1). (B-D) Western blot analysis of the distribution of SorCS2 and EAAT3 in subcellular brain fractions from wild-type (S2+) and S2KO mice (S2−). SorCS2 and EAAT3 are highly enriched in synaptosomal fractions (synaptos.; B), light membranes fraction (light m.; C), and extra-synaptic membranes fraction (extras. m.; D). The efficiency of brain fractionation is documented by detection of specific marker proteins: Rab11 (light membranes), synaptophysin (synaptosomes), and GluA1, GluA2, and PSD95 (PSD1 fraction). Actin served as loading control. (E) Representative images showing immunodetection of SorCS2 (red, upper panels) and EAAT3 (red, lower panels) with markers of recycling endosome (Rab11, green) and early endosome (Rab5, green) in primary wild-type neurons. Cell were counterstained with DAPI. Scale bar: 10 μm. Insets show higher magnification of the given images.

**Figure S3. Related to Figure 5.**
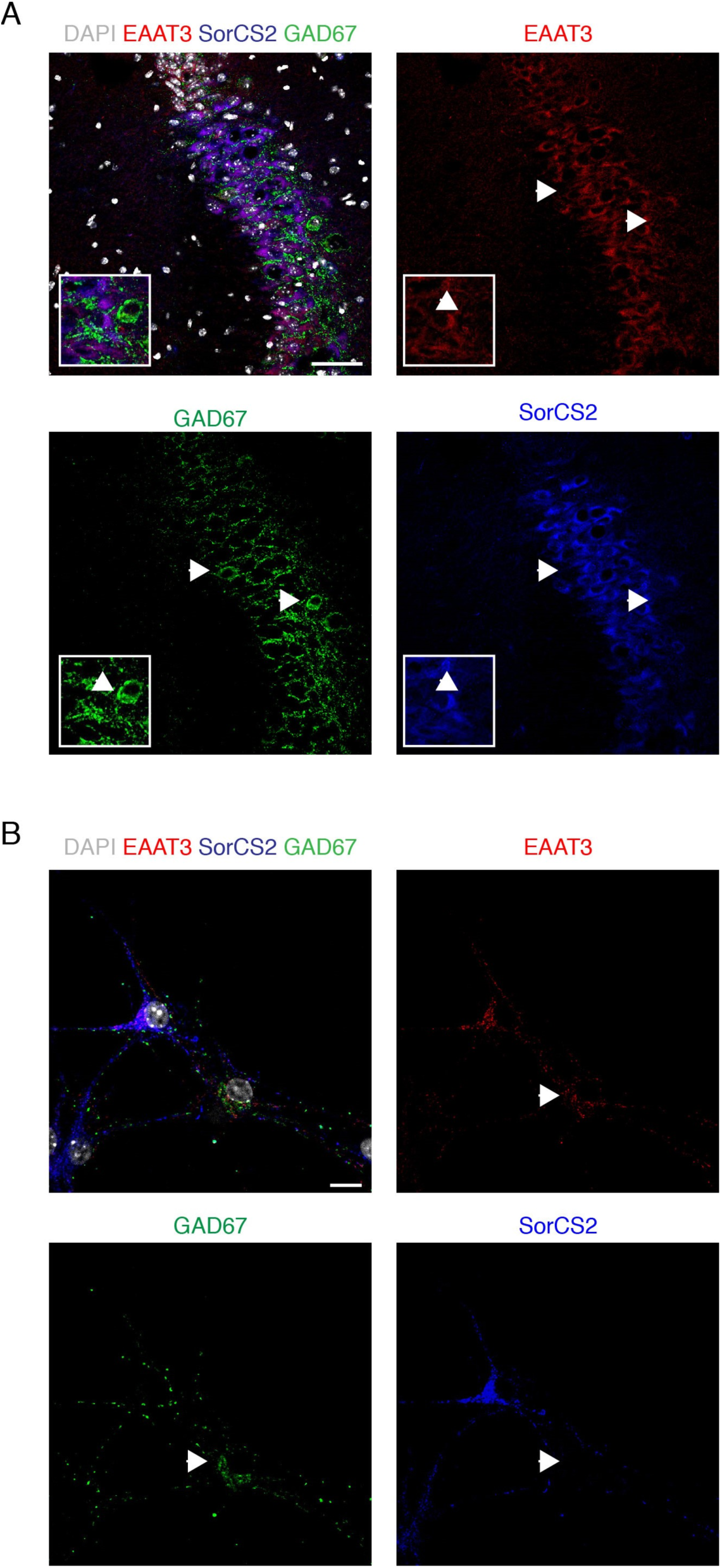
SorCS2 is not expressed in GAD67-positive inhibitory neurons. (A) Representative confocal images showing immunodetection of SorCS2 (blue), EAAT3 (red), and the GABA-ergic neurons marker GAD67 (green) in the CA2 region of the mouse hippocampus. Cells were counterstained with DAPI (white). Both individual channels and merged immunostainings are shown. SorCS2 is not expressed in GAD67-positive neurons (indicated by arrowheads). Scale bar: 50 μm. (B) Representative images showing immunodetection of SorCS2 (blue), EAAT3 (red), and the GABA-ergic neurons marker GAD67 (green) in cultured hippocampal neurons. EAAT3 is detected in GAD67-positive neurons (indicated by arrowheads), but SorCS2 is only expressed in GAD67-negative neurons. Scale bar: 10 μm.

**Figure S4. Related to Figure 5.**
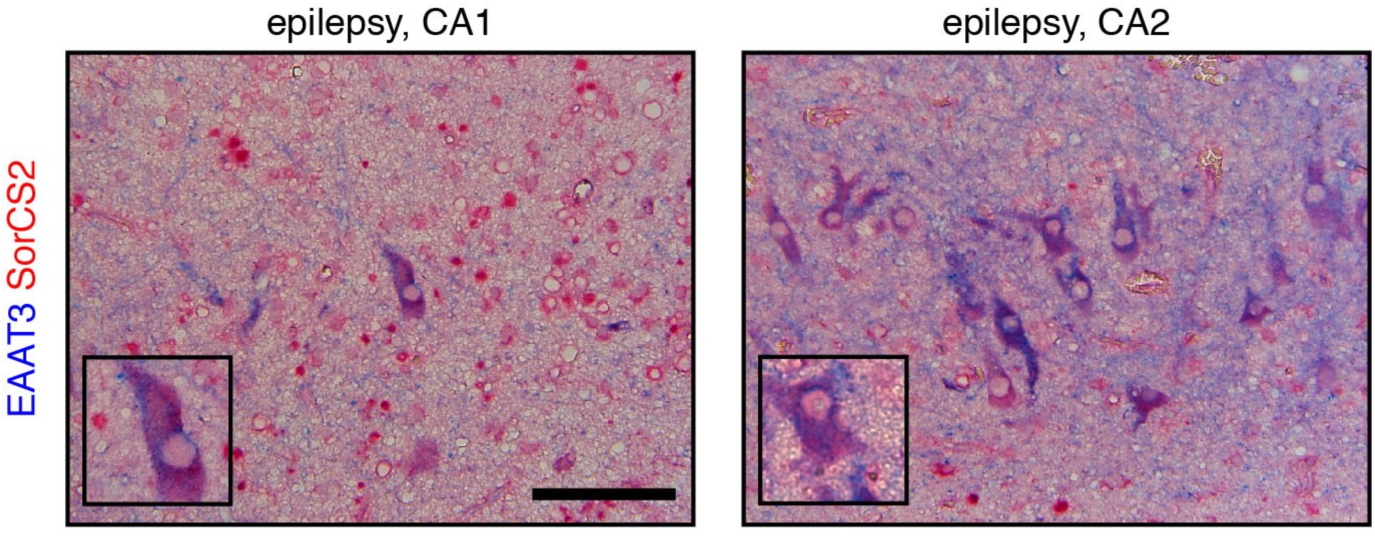
SorCS2 and EAAT3 colocalize in neurons of the human epileptic hippocampus. Representative images showing immunodetection of SorCS2 (red) and EAAT3 (blue) in surviving neurons of the CA1 and CA2 regions of the human epileptic hippocampus. Insets show higher magnification of the given images. Scale bar: 100 μm.

